# Environmental and molecular modulation of motor individuality in larval zebrafish

**DOI:** 10.1101/2021.09.14.460248

**Authors:** John Hageter, Matthew Waalkes, Jacob Starkey, Haylee Copeland, Heather Price, Logan Bays, Casey Showman, Sean Laverty, Sadie A. Bergeron, Eric J. Horstick

**Author notes:** Correspondence: Eric Horstick.

## Abstract

Innate behavioral biases such as human handedness are a ubiquitous form of inter-individual variation that are not strictly hardwired into the genome and are influenced by diverse internal and external cues. Yet, genetic and environmental factors modulating behavioral variation remain poorly understood, especially in vertebrates. To identify genetic and environmental factors that influence behavioral variation, we take advantage of larval zebrafish light-search behavior. During light-search, individuals preferentially turn in leftward or rightward loops, in which directional bias is sustained and non-heritable, and maintained by a habenula-rostral PT circuit. Here we use a medium-throughput recording strategy and unbiased analysis to show that significant individual to individual variation exists in wildtype larval zebrafish turning preference. We classify stable left, right, and unbiased turning types, with most individuals exhibiting a directional preference. Raising larvae at elevated temperature selectively reduces the leftward turning type and impacts rostral PT neurons, specifically. Exposure to conspecifics, variable salinity, environmental enrichment, and physical disturbance does not significantly impact inter-individual turning bias. Pharmacological manipulation of Notch signaling and carrying a mutant allele of a known Notch pathway affecter gene, *gsx2*, disrupted turn bias individuality in a dose-dependent manner. These results establish that larval zebrafish is a powerful vertebrate model for inter-individual variation with sensitivity to specific environmental perturbations and gene dosage.

## Introduction

Inter-individual variation, or individuality, is a hallmark of nearly all animal species and contributes to the population’s fitness and ability to adapt when confronted with environmental change (Dingemanse et al., 2004; Klein et al., 2017; Horváth et al., 2020). One form of inter-individual variation is sensory and motor biases. Handedness in humans is a familiar example, expressed as a sustained preference for left- or right-hand use, which the expression of a specific hand preference is only partially explained by genetics, suggesting complex interactions contribute to imposing handed phenotypes (Cuellar-Partida et al., 2020). Indeed, significant complexity underlies human handedness. The preferred hand usage is context-dependent, largely independent of other behavioral biases, and shows variable consistency – observed as consistent or inconsistent hand dominance in a task dependent manner (Watson and Kimura, 1989; Souman et al., 2009; Chu et al., 2012). Handed biases are also a conserved form of individual behavioral variation with species as diverse as hagfish (Miyashita and Palmer, 2014), *Drosophila* (Kain et al., 2012; Buchanan et al., 2015), chicken (Rogers, 1982; Casey and Karpinski, 1999), and various vertebrate paw/foot biases (Bulman-Fleming et al., 1997; Brown and Magat, 2011; Giljov et al., 2013; Schiffner and Srinivasan, 2013; Manns et al., 2021) showing sustained individual motor preferences. Despite the prevalence of handed behaviors, mechanisms instructing these behaviors and the variation observed across individuals are still poorly understood.

Two prevailing questions are what neural substrates generate biases and what mechanisms instruct specific bias types, i.e., left versus right-handed or consistent versus inconsistent handedness. Research to date shows that isogenic *Drosophila* (Kain et al., 2012; Buchanan et al., 2015; Linneweber et al., 2020), *C. elegans* (Stern et al., 2017), mouse strains (Freund et al., 2013, 2015; Hager et al., 2014), clonal fish (Izvekov et al., 2012; Bierbach et al., 2017), and clonal crayfish (Vogt et al., 2008) species display stable individual phenotypes with significant inter-individual variation at the population level, suggesting external events contribute to behavioral diversity across individuals. Even in humans, external or stochastic factors are likely important as discordant handedness is frequently observed in monozygotic twins (Jäncke and Steinmetz, 1995). In *Drosophila*, the availability of numerous isogenic strains and the ability to assay large numbers of individuals have been instrumental in elucidating key components generating inter-individual variation (Buchanan et al., 2015). When navigating in their environment, *Drosophila* display a turn bias, where individuals preferentially use same-direction turns, and the magnitude of this bias is modulated by genetic background, activity in the central complex, and exposure to environmental enrichment as well as social experiences (Ayroles et al., 2015; Buchanan et al., 2015; Akhund-Zade et al., 2019; Versace et al., 2020). These findings demonstrate that functional variation in the invertebrate nervous system is maintained by specific neural substrates and further modified by gene and environment interaction. In murine models, exploratory behavior is a thoroughly investigated example of inter-individual variation, where phenotype variation is enhanced by environmental enrichment (Freund et al., 2013; Körholz et al., 2018; Zocher et al., 2020). Despite this well-studied mammalian model and other known handed behaviors, the regulation of inter-individual variation in vertebrates remains poorly understood.

Zebrafish have emerged as a powerful model for elucidating mechanisms that instruct visceral and neural differences between individuals (Gamse et al., 2003, 2005; Dreosti et al., 2014). Moreover, similar to other teleost species, zebrafish have a visual bias, preferentially using the left eye to assess novelty (Bisazza et al., 1997; De Santi et al., 2001; Sovrano, 2004; Andrew et al., 2009). However, this behavioral bias is primarily fixed in the population and offers little insight into inter-individual variation. Larval zebrafish also perform a light-search behavior that is onset by the loss of visual navigating cues, which is characterized by a period of stereotypic leftward or rightward circling (Horstick et al., 2017), consistent with search patterns observed in other species (Bell et al., 1985; Hills et al., 2004, 2013; Gray et al., 2005). An individual’s leftward or rightward circling direction is persistent over multiple days, observed at equal proportions in the population, and is not heritable (Horstick et al., 2020). The features of light-search share many of the hallmark traits observed in well-established invertebrate models of turn bias that have been instrumental for characterizing mechanisms that regulate inter-individual variation (Ayroles et al., 2015; Buchanan et al., 2015; Akhund-Zade et al., 2019). Moreover, our work has shown that neurons in the habenula and rostral posterior tuberculum (PT) are essential for maintaining zebrafish turn bias (Horstick et al., 2020). Therefore, larval zebrafish is a potentially powerful vertebrate model to determine how inter-individual variation is imposed in the vertebrate brain. What remains lacking is a rigorous analysis of turn bias variation in the population and the identification of external and internal factors modulating inter-individual turn bias differences.

Here, we capitalize on the larval zebrafish turning bias to describe environmental factors and signaling pathways that modulate inter-individual variation. First, we develop a multiplex recording pipeline to record behavior in a medium-throughput manner, allowing a rigorous analysis of the spectrum of inter-individual variation. Using this pipeline, we characterized stable biased and unbiased turning types in a wildtype strain, distinguishable by unique path trajectory features. Second, we determined that temperature fluctuation impacts inter-individual variation during early development, yet this is unaffected by conspecifics, environmental enrichment, and stressful stimuli. Last, pharmacological and mutant analysis shows that maintaining turn bias shows dose sensitivity to signaling pathways associated with neuronal proliferation.

## Methods

### Animal Husbandry

All experiments were approved by the West Virginia University Institutional Animal Care and Use Committee. Zebrafish (*Danio rerio*) Tubingen long-fin (TL) wildtype strain was used in all experiments and used as the genetic background to maintain transgenic and mutant lines. Experiments were conducted during the first 7 days post fertilization (dpf), which is before sex determination. Larval rearing conditions were 28°C, 14/10 hour light-dark cycle, in E3h media (5 mM NaCl, 0.17 mM KCl, 0.33 mM CaCl2, 0.33 mM MgSO4, and 1 mM HEPES, pH 7.3), and at a stocking density of 40 embryos per 30 mL E3h, unless stated otherwise. *Social environment*: To test the effect of social interaction, we raised larvae under two different social conditions: 20 larvae in a 6 cm petri dish or a single larva per 6cm dish. Social or isolation rearing started at 5-8 hours post fertilization (hpf) and continued until testing at 6 dpf. *Temperature*: To test the impact of temperature on the development of turn bias, larvae were raised 1-4 dpf at either 24, 28, or 32°C. At 4 dpf, all groups were moved to 28°C until testing at 6 dpf. To determine if a specific development period was sensitive to elevated temperature, separate groups of larvae were raised at 32°C from either 31-55 hpf or 55-79 hpf, after which they were returned to standard rearing temperature and tested at 6 dpf. *Salinity*: The impact of increased salinity was tested over 4 salt concentrations (1 ppt, 2ppt, 5ppt – parts per thousand) and standard E3h (∼0.5 ppt) as a control. Larvae were reared in variable salinities from 1-4 dpf, and behavior tested at 6 dpf. An elevated salinity stock of E3h was made by adding 9.5g NaCl (Sigma) to standard E3h, creating a 10 ppt stock, which was diluted for working concentrations with standard E3h media. *Environmental enrichment:* Enriched environments were created by adhering mixed size and color (predominately red, blue, grey, and white colors) LEGO® blocks onto the bottom of a 10 cm petri dish. Previously, LEGO® blocks have been used to stimulate novel object recognition and interaction in larval zebrafish (Bruzzone et al., 2020). In addition, 5-8 plastic aquarium leaves were included to float on the surface. Last, dishes were positioned on platforms with mixed white and black shape substrates. A total of four enriched environments were created with variable LEGO® colors and sized blocks, and larvae were rotated daily between enriched environments. As controls, larvae were raised in plain 10 cm dishes placed on either a solid white substrate. For experiments, larvae were maintained in enriched or control dishes from 1 dpf until behavior testing. *Shaking*: We tested the impact of environmental instability on motor bias by continuously shaking larvae from 1-4 dpf. At 1 dpf, embryos were placed in a 75 cm^2^ cell culture flask (Sigma) with approximately 80 mL E3h. Flasks were propped at 30 degrees on a Stovall Belly Dancer orbital rotator, set to 70 rpm. At 4 dpf, larvae were removed from culture dishes and raised under standard conditions prior to testing at 6-7 dpf.

Transgenic lines used were enhancer trap *Tg(y279-Gal4)* (Marquart et al., 2015) and *Tg(UAS:Kaede)s1999t* (Davison et al., 2007). Mutant line used was *gsx2*^*Δ13a*^ (Coltogirone et al., 2021).

### Behavior tracking and analysis

Behavioral experiments were performed on 6-7 dpf larvae, except as noted. All experiments were recorded using infrared illumination (940nm, CMVision Supplies), a µEye IDS1545LM-M CMOS camera (1^st^ Vision) fitted with a 12mm lens, and a long-pass 780 nm filter (Thorlabs, MVL12WA and FGL780, respectively). Visible illumination was provided by a white light LED (Thorlabs) positioned above the larvae, adjusted to 40-50 µW/cm2 (International Light Technologies, ILT2400 Radiometer with SED033 detector). Testing conditions were maintained between 26-28°C for all behavioral recording, and all larvae adapted to the recording room conditions for 20 minutes before recording under matched illumination to recording rigs. Custom DAQtimer software was used to control lighting, camera recording parameters, and real-time tracking as previously described (Yokogawa et al., 2012; Horstick et al., 2017). The camera field of view was set to record four 10 cm dishes simultaneously with one larva per dish for multiplex recordings. A total of four recording rigs were used. Path trajectories of individual larvae are recorded over 30-second recording intervals at 10 fps and analyzed using five measures: net turn angle (NTA), total turning angle (TTA), match index (MI), bias ratio (BR), and performance index (PI). A minimum of 100 points were required to be included in the analysis. NTA is the summation of leftward and rightward angular displacement (-leftward, + rightward) over the recording interval, whereas TTA is the sum of absolute values of all angular displacement. MI measures the proportion of events in a series going in the same direction. Leftward and rightward trials are scored as 0 or 1, and MI is the percent of events matching the direction of the first trial in a testing series. For example, a MI=1 is all events are in the same direction as the first trial, whereas 0.33 is a third of the events matching the first trial. For MI analysis, individuals missing the first dark trial were excluded from analysis. BR is a proportion of directional turning compared to total turning, calculated by dividing NTA by TTA, e.g., -1 represents that all directional movement in a single trial occurred in a leftward direction, while -0.5 indicates that half of all directional movement was in a net leftward direction. PI was calculated by averaging binary bias ratios, with leftward trials scored as 0 and rightward 1. Bias ratios were weighted by the proportion of larvae within a PI group. In all analyses that required a PI for categorizing larvae, whether in 4X, 8X, or q4X recordings, all individuals that had missing trials were excluded. This criteria was necessary to ensure rigorous categorization. For *gsx2* experiments, larvae were housed individually following behavior testing for posthoc genotyping. Genotyping was performed as previously described (Coltogirone et al., 2021). In brief, genotypes were confirmed using PCR spanning the deletion: *gsx2* (primers: 5’TGCGTATCCTCACACATCCA, 5’TGTCCAGGGTGCGCTAAC; 134 bp wildtype, 121 bp mutant, and 134/121 bp heterozygous). Previous reports describe that gsx2 mutants have reduced swim bladder inflation (Coltogirone et al., 2021), which was minimized by raising larvae in shallow water dishes. Only larvae with normal swim bladder inflation and balance were used for experiments.

The 4X recording assay was performed by recording larval path trajectories over four recording intervals, each composed of 30 seconds baseline recordings, immediately followed by 30 seconds recording following the loss of visible illumination. Each recording interval was separated by three minutes of baseline illumination. The 8X recording was performed in a similar format, including four additional light ON/OFF recording intervals performed in series. The quad 4x (q4X) assay is identical to the 4X, except that the 4X recording interval is repeated four times, separated by 10 minutes baseline illumination. A 4X recording strategy was used to test the developmental onset of turn bias. Individual larvae were first tested at 3 dpf, and were separately raised in 6-well plates and retested daily through 6 dpf. For analysis, larvae were grouped as left or right biased based on NTA (average NTA +, right bias; -, left bias) at 6 dpf when turn bias is well-established (Horstick et al., 2020).

### Pharmacology

Notch signaling was inhibited using the γ-secretase inhibitor LY411575 (Sigma, SML050). A 10mM stock of LY411575 was prepared in DMSO and diluted to working concentrations with a final volume of 0.08-0.1% DMSO for all trials. To test Notch inhibition on turn bias, mid-gastrulation (6-8 hpf) groups of embryos were treated with 0.05, 0.1, 0.15, 0.2, 0.25, 1 or 10 µM LY411575 until 4 dpf; the drug was replaced daily. At 4 dpf, LY411575 was removed and larvae placed in fresh E3h until behavior testing at 6 dpf. Phenotypic categorization was performed at 3 dpf. Individuals were scored as normal (no morphological or behavioral defects), mild (abnormal touch responsiveness), moderate (tail curvature), or severe (gross developmental defects, necrosis). Only normal larvae were used for behavioral testing. Vehicle controls were 0.08-0.1% DMSO treated.

### Labeling and imaging

#### Immunohistochemistry

To assay neuronal proliferation, we labeled with anti-HuC/D (Elav protein) (Invitrogen A21271). Control (0.08% DMSO) and LY411575 groups (100 nM and 8 µM) were prepared as described above. At 24 hpf, embryos were fixed overnight using 4% paraformaldehyde in 1X PBS at 4°C. Washes were performed with 1X PBS containing 0.1% TritonX-100. We used primary antibody mouse anti-HuC/D (1:500, Invitrogen, 16A11). Secondary detection was performed with goat anti-mouse IgG1 Alexa 488 (1:500, Invitrogen, A32723). To analyze images, signal intensity of a 56 × 6 µM (W x H) region spanning a lateral to midline hemi-section of the anterior spinal cord was recorded using ImageJ. Three sections were measured per larva, averaged and standardized for comparison between groups.

#### Fluorescent in-situ hybridization

To determine the levels of Notch signaling we examined transcript levels of her12 (Jacobs and Huang, 2019). Hybridization chain reaction (Molecular Instruments) probes and labeling technology was used to detect her12 transcripts. Her12 mRNA sequence (NM_205619) was provided to Molecular Instruments to design a custom gene-specific her12 probe detection set. LY411575 and control larvae were treated as described above. At 30 hpf, larvae were fixed overnight using 4% paraformaldehyde in 1X PBS at 4°C. Fixed larvae were washed in 1X PBS containing 0.1% Tween20 and labeled following Molecular Instruments HCR RNA-Fish protocol for whole-mount zebrafish embryos (Schwarzkopf et al., 2021). All images were collected using the same parameters. For analysis, the percent area of her12 expression was quantified within the spinal cord using ImageJ.

#### Neuron temperature sensitivity

Rostral PT and habenula neuron areas were quantified using larvae from *Tg(y279-Gal4)/Tg(UAS:kaede*) carrier in-crosses. At 1 dpf, larvae were screened for Kaede and reared at elevated temperatures as described above. Larvae were moved to standard raising conditions at 4 dpf, and live-imaged at 6 dpf. Larvae were anesthetized using MS-222 (Sigma) and mounted in 2% low melting temp agar. Image stacks were collected through the rostral PT and habenula for experimental and control groups and area quantified for analysis in ImageJ. To determine if a specific developmental time period was crucial, larvae were similarly prepared and analyzed, yet only raised at elevated temperature during either 31-55 hpf or 55-79 hpf intervals. Controls were raised at standard rearing temperatures.

#### Imaging

All imaging was performed on an Olympus Fluoview FV1000. For live imaging, larvae were anesthetized in a low dose of MS222 (Sigma) and embedded in 2% low melting temp agar. Fixed samples were transferred into 70% glycerol/30% 1X PBS and slide-mounted for imaging.

### Statistical analysis

Analysis was performed in R (R Core Team (2020). R: A language and environment for statistical computing, 2020), R ggplot2 package (Wickham, 2016) (R Core Team) and Prism (GraphPad). All statistical comparisons were two-sided, unless noted otherwise. Standard error of the mean (± SEM) was used for all experiments, except MAD analysis which display 95% confidence intervals. Cohen D was calculated in R using package effsize. For all experiments, data was collected from a minimum of three independent biological groups. Normality was tested using the Shapiro-Wilks test. Normally distributed data was compared using either 1 or 2-way *t*-tests. Non-normal data was analyzed using a Mann-Whitney U test or Wilcoxon signed-rank test for 2 or 1-way tests, respectively. To perform multiple comparisons, ANOVAs were performed in GraphPad and multiple comparisons adjusted using a Bonferroni correction. Boxplots show median and quartiles with outliers identified beyond 2.7 standard deviations from the mean.

Reshuffling and bootstrapping was performed using “sample” R function without and with replacement, respectively. For reshuffling experiments, bias ratios values were randomized across all individuals in a dataset. Randomization was performed only within the same trial, e.g. reshuffling of bias ratios within the first light off trial. Randomization was simulated 1000 times and average reshuffled bias ratios and MAD values calculated using custom R code, and used to plot reshuffled probability density curves and reshuffled MAD values. Probability density plots and area under the curve measurements were performed using custom R code. For area under the curve analysis, ±0.3 tails were chosen for comparison, which are approximately two standard deviations from the population average. To generate error bars for MAD analysis, average bias ratios were bootstrapped (1000 bootstrap replicates) with resampling. For each resampled dataset a MAD was calculated and MAD values across all resampled datasets used to calculate a 95% confidence interval applied as an error bar. A 1-way comparison was used to calculate significance for all simulated dataset comparisons. To generate a p-value, the number of resampled dataset MAD values were totaled that fall within or exceed the 95% confidence interval of the comparison group, and this total was divided by 1000 to produce a final p-value. This represents the fraction of simulated experimental groups that fall within a range that supports a null hypothesis of no difference between groups. For example, 600 bootstrapped datasets from a simulated control that fall within or exceed the confidence interval of an experimental group yields p=0.60, implicating that 60% of simulated datasets do not support the statistical difference between compared groups. Direction of comparison is noted in the legend for each dataset.

## Results

### Turning behavior during light search shows high inter-individual variation

We developed a multiplexed strategy to record path trajectories to assess inter-individual variation during larval zebrafish light search behavior (Figure 1A). Previously, the stereotypic turning was described using a large recording arena (14400 mm^2^) to record single larva (Horstick et al., 2017). Larvae are recorded in 100 mm diameter dishes (7854 mm^2^) for our multiplexed strategy, and robust circling is observed following light extinction (Figure 1B). To characterize individual motor biases, we initially recorded larval path trajectories over a series of four intervals of paired 30-second baseline illumination and 30 seconds following the loss of illumination, with each of these recording pairs separated by 3 minutes of illumination to restore baseline behavior (Horstick et al., 2020), which we refer to as 4X recording (Figure 1C). This recording yields four paired light on and off events per individual. We recorded responses from 374 individuals, representing 1496 paired baseline and dark responses. The presence of motor bias was previously described using a match index (MI) - the percent of turning trials in which turning direction was the same as the first dark trial (Horstick et al., 2020). Here we confirm previous findings showing a significant MI increase following the loss of illumination (Wilcoxon matched-pairs test, p<0.001), which can be scaled via multiplexing (Supplementary Figure 1A). Overall, our current approach for multiplexed recording recapitulates previous findings. These data show that our multiplexed strategy provides medium-throughput recording, allowing a rigorous analysis of larval zebrafish inter-individual variation.

**Figure 1.**
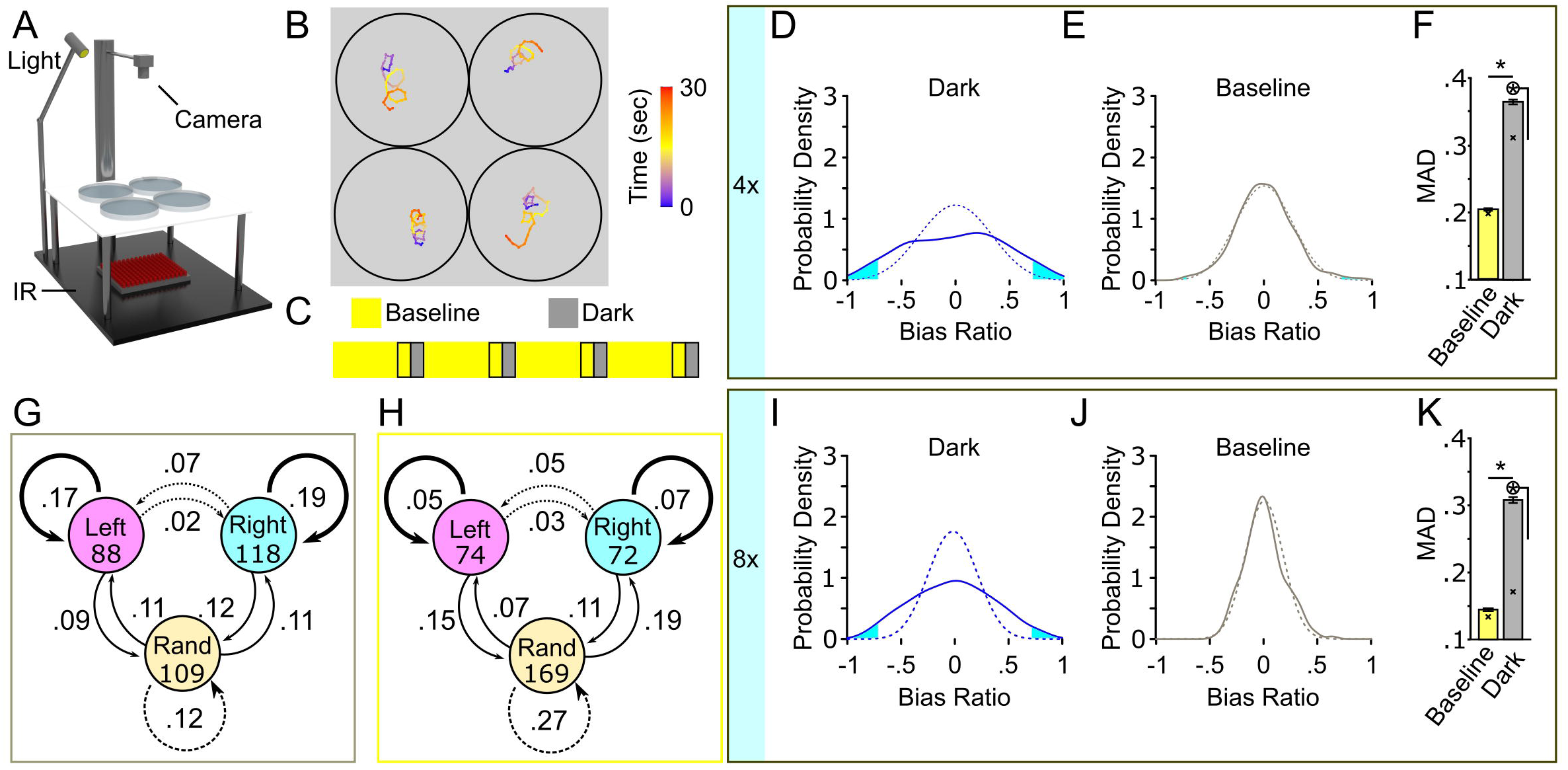
Turning behavior following the loss of light exhibits high inter-individual variation. **(A)** Schematic of multiplex recording rig. **(B)** Illustrative path trajectories following loss of light from multiplex recording. Color scale is time (seconds). **(C)** Diagram of 4X recording. Outlined regions denote recording intervals. **(D-F)** 4X recordings strategy to show high inter-individual variation in turn bias. **(D)** Average bias ratio probability density curve for baseline responses (solid grey line, N=374) and paired curve following reshuffling (dotted grey line, average of 1000 resampled datasets). **(E)** Same as C except the paired dark responses following the loss of illumination (solid blue line, N=374) and reshuffled density curve (dotted blue line). Cyan fill shows the area in each tail corresponding to the probability of observing a result more extreme or equal to ± 0.3 average bias ratio. **(F)** MAD for baseline (yellow), and dark (grey) responses. N=374. Error bars are 95% confidence intervals generated by bootstrap resampling. Asterisk in circle, p < 0.05 to reshuffled MAD value shown with an X. **(G)** Dark responses (grey outline) and baseline **(H)** (yellow outline) from 4X recording showing frequency of direction change between first (circles) and second (arrows) pairs of responses. Initial N shown in circles. Bold, solid, and dotted arrows delineate responses that produce perfectly matched bias, partial bias, and unbiased responses, respectively. **(I-K)** Same analysis as in C-E except for 8X recordings. Asterisk p <0.05. Asterisk in circle, p <0.05 to reshuffled MAD value.

We calculated a bias ratio by dividing net turning angle (NTA) by total turning angle (TTA – absolute sum of all angular displacement) for each baseline, and dark trial recorded to examine the spectrum of wildtype larvae inter-individual variation during search behavior (Supplementary Figure 1B). This metric provides the proportion of same-direction turning within a continuous numerical range bounded by -1 and 1, representing all directional movement in a leftward or rightward direction, respectively. The average bias ratio across the entire population during baseline illumination and light-search did not significantly vary from zero showing no population bias (one-sample *t*-test against 0, baseline: *t(373)* = 0.007842, p=0.9937; dark: *t(373)* = 0.1696, p=0.89)(Supplementary Figure 1C). Despite similar population-level bias ratios between baseline and dark, significant variation is observed in the dark that is not observed during baseline (Figures 1D-F). Using a probability density curve, where the area under the curve represents the proportion of individuals in the population, we find that during dark turning, 12.38% of the population displayed a robust sustained turning bias over 4 trials (bias ratio <-0.7 = 6.41%, left bias; >0.7 = 5.97%, right bias) (Figure 1D). Conversely, 1.72% of baseline events displayed sustained directional turning (Figure 1E). The distribution of bias ratios shows that, following light extinction, a significantly greater number of individuals utilize sustained same-direction turning (χ^2^(1) = 51.02, p<0.0001). To determine whether these distributions were the product of chance, we simulated ‘randomized’ baseline and dark datasets by resampling bias ratios (1000 resamples) within each trial (Figures D-E, dotted line). Following randomizing, 2.35% of the simulated dark responses maintained strong directional turning, similar to that observed during baseline. A previous study used mean absolute deviation (MAD) as a metric to quantify variation in a population; a higher MAD represents increased variation across individuals in the population (Buchanan et al., 2015). Here, MAD was calculated for baseline, dark, and simulated datasets. As MAD was generated from the whole population, average bias ratios were bootstrapped (1000 boots) to generate 95% confidence intervals for statistical comparison. MAD is 44.10% (p<0.001) and 15.79% (p<0.001) reduced in baseline or in randomized dark groups compared to light-search dark responses, respectively (Figure 1F), whereas no difference was observed between baseline MAD and randomized baseline responses (Figure 1F, yellow bar). These findings show that turn bias during light search behavior shows significant variation beyond what is expected by chance or while larvae navigate in an illuminated environment.

Our analysis, along with findings from previous reports, illustrates robust left and right turners, or turning types, within the population. However, the distribution of bias ratios from 4X recordings shows that over 14% of the population exhibits an average bias ratio consistent with no sustained turn direction (−0.1 < BR < 0.1) (see Figure 1D). These individuals could represent either a stable unbiased population or endogenous behavioral fluctuation. To evaluate whether unbiased individuals are a sustained turning type in the population, in addition to left/right biased turners, we created a performance index (PI) by transforming all individual trials to either 0 or 1 for overall leftward or rightward preference per trial, respectively. From these binary values, we created a transition index for the first and second set of responses from the 4x dataset, i.e. left (LL = 0), right (RR = 1), or random (LR; RL = 0.5) responses that can be compared between the first and last response pairs. Using the transition pair PI, we assessed the frequency of turn direction change or conservation (Figure 1G-H). During dark trials, 36% of all transitions showed sustained turn direction (left = 17%, right = 19%; average PI = 0 or 1), whereas during baseline illumination 12% of larvae sustained turn direction (χ^2^(1) = 54.545, p<0.0001). Conversely, 21% and 35% of transitions yielded sustained random behavior between dark and baseline recording conditions, respectively (for example, LR to RL or RR to LL; average PI = 0.5) (χ^2^(1) = 8.615, p=0.0033) (Figure 1G-H). Interestingly, during light-search initially random response pairs transitioned to directional (RR or LL) responses 22% of the time, yielding partial turn bias (average PI 0.75 or 0.25).

To confirm our observations persisted over longer timescales, we ran an additional 8X experiment, testing 189 larvae as before, with four additional light ON/OFF intervals in series. From this extended testing condition, we observed conserved trends demonstrating significant inter-individual variation in turning bias during light-search, yet not during baseline illumination (Figure I-K). Neither 4X or 8X recording showed a change in TTA over time, establishing overall behavior is not disrupted by our assays (Supplementary Figure 1D-E). As 4x and 8x experiments were broadly consistent, we focused on the 4X recording strategy for ongoing investigations. Our data show that wildtype larvae exhibit significant inter-individual variation in turn bias during light-search, greater than that expected by chance, with a subset of individuals potentially exhibiting a previously unexplored unbiased turning type.

### Multiple stable turning types exist with distinct locomotor features

Characterizing changes in locomotor parameters in zebrafish has been a powerful strategy to develop etiological and mechanistic models (Burgess and Granato, 2007; Horstick et al., 2013; Chen and Engert, 2014; Dunn et al., 2016). Therefore, we next aimed to establish what underlying locomotor changes account for unbiased and biased motor types. We hypothesized that three possible modes could generate unbiased behavior: 1) normal turning with high rates of direction switching across trials, 2) reduced same-direction turning within single trials, or 3) weak photo-responsiveness and, therefore, low total turning. To differentiate between these hypotheses, we categorized all larvae based on average PI across all four trials, generating five categories. Across PI groups, we compared the absolute average bias ratio to determine if the magnitude of directional turning changed based on PI. During light search the average bias ratio magnitude significantly changed based on PI (1-way ANOVA *F(4,352) = 10*.*43*, p<0.0001), where partial and unbiased PI groups showed less overall directional turning (Figure 2A). No difference was observed between strong left and right biased turners (PI =0, 0.603 ± 0.022; PI = 1, 0.58 ± 0.021: *t(352)* = 0.7811 adjusted p>0.9999). Consistent with earlier observations, no differences were observed across PI groups during baseline (1-way ANOVA *F(4,352) = 2*.*087*, p=0.082), consistent with an absence of turn individuality (Supplementary Figure 2A). Moreover, there was no significant change in TTA during dark turning (1-way ANOVA *F(4,352) = 1*.*263*, p=0.28) across all PI groups (Supplementary Figure 2B). As all PI groups exhibited normal levels of total turning, this ruled out variable photo responsiveness as the basis of different turning types. Unexpectedly, partially biased populations (0.25, 0.75 PI) showed a similar average bias ratio as unbiased larvae (Figure 2A). To explain this observation, we reasoned that bias ratio magnitude might vary depending on whether an individual trial matches or opposes the overall larva turning type. For example, for 0.25 PI larvae, leftward matched direction bias ratios compared to rightward opposed direction trials. We analyzed all individual trials between all performance groups to explore this idea, sorting trials into matched or opposing based on the average PI for each individual. Perfect performance trials (0,1) were categorized as all matched, whereas unbiased trials (0.5) as all unmatched. Left and right direction bias ratios did not vary in these groups; therefore, we combined these groups to simplify comparison (Supplementary Figure 2C). A significant effect was observed across groups (1-way ANOVA *F(3,1408) = 27*.*93*, p<0.0001), with trials opposed to overall PI direction showing lower overall bias ratio strength (Figure 2B, magenta lines). These data suggest that the basis of unbiased motor types is due to a lower bias ratio or less persistent same-direction turning, yet not a loss of overall turning. Interestingly, we noted that matched bias ratios were reduced in partially matched trials compared to events in the fully matched group (match 0.594 ± 0.013: partial match 0.514 ± 0.015: *t(1408)* = 4.046 adjusted p=0.003)(black line), implicating that the underlying differences between biased and unbiased larvae may be graded.

**Figure 2.**
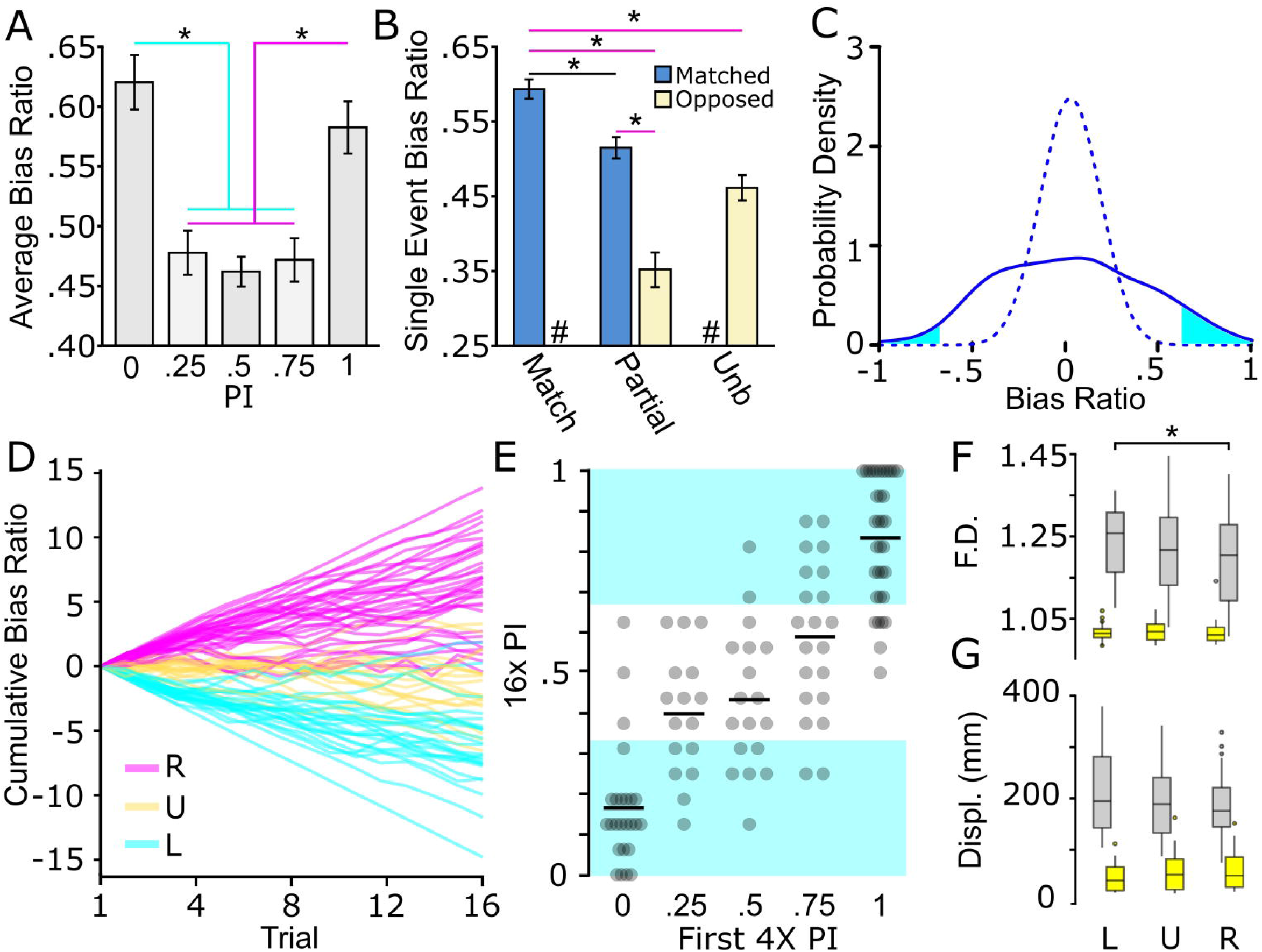
Biased and unbiased motor types present during light-search. **(A)** Absolute bias ratio from 4X recording per PI (PI 0, N=66; PI 0.25, N=74; PI 0.5, N=75; PI 0.75, N=75; PI 1, N=67). **(B)** Absolute bias ratio magnitude for single events occurring in a direction matching (blue) overall average PI direction or occurring in the opposed (beige) direction shown for perfect bias (0,1 PI, N=517), partially biased (0.25,0.75 PI, N=596), and unbiased (05 PI, N=301) populations. **(C)** Average q4X probability density distribution for dark responses (solid line, N= 114) and reshuffled populations generated from the average of 1000 resampled populations (dotted line). Cyan fill displays percent of population with strong left (<-0.3) and right (>0.3) average bias ratios. **(D)** Cumulative summation of bias ratio over the q4x from strong right (magenta, N= 34), strong left (cyan, N= 34, and unbiased (yellow, N= 18) individuals. Individuals were selected based on the first 4X average PI (strong right PI=1, strong left PI=0, unbiased PI=0.5). (**E)** Average q4x PI based on initial 4x PI (PI=0, N=66; PI=0.25, N=74; PI=0.5, N=75; PI=0.75, N=75; PI=1, N=67). Black bars represent sample mean. Left, right, and unbiased performance categorized within equal thirds of the PI scale. Cyan background highlights average PI with left or right turning type behavior. **(F-G)** Characterization of path trajectory features during baseline (yellow boxes) and dark (grey boxes) responses from individuals tested in the q4X assay. Turning type determined by 16X average PI distribution in E. (**F)** Fractal dimension and **(G)** Displacement (L=left, N=36; U=unbiased, N=40; R=right, N=38). Asterisk p <0.05.

In order to confirm rigorously the three motor types, we performed a quad 4X assay (q4X), using the standard 4X assay repeated four times, with each recording sequence separated by 10 minutes of baseline illumination (Supplementary Figure 2D). We recorded 114 larvae, and consistent with our previous measures, individuals showed significant inter-individual turn bias variation during light-search (± 0.3 probability density tails: 7.34% dark; 0.00037%, randomized dark), and sustained left, right, or unbiased locomotor preferences (Figure 2C-D). The cumulative summation of bias ratios provided a qualitative measure of turn performance over time (Figure 2D). From this analysis, we noted that some larvae initially categorized as strong or unbiased turners, seemingly switched over time. Therefore, we next aimed to utilize the q4X analysis to quantify bias determination accuracy by comparing the first 4X PI to overall q4X performance. We equally divided the 0 to 1 PI scale for classifying left, right, or unbiased behavior (left ≤ 0.33; unbiased 0.33 < 0.66; right ≥ 0.66) (Figure 2E). Of the larvae that show an initial strong or partial bias during the first 4x interval, 2/96 (2.08%) reverse bias direction during the q4x assay, and 27/96 (28.13%) of these individuals ultimately switch to an unbiased response after serial q4x testing. However, switching is primarily observed in larvae showing an initially partial bias, as the larvae that displayed an initially strong bias (0, 1 PI) in the q4x assay, 50/59 (84.75%) maintained a left or rightward turning type. Interestingly, at the population level, the 9/114 (7.89%) of unbiased individuals initially categorized with a strong bias was comparable to that expected by random chance, i.e., the same 6.25% likelihood of flipping 4 heads with a coin (χ^2^(1) = 0.609, p=0.435). As expected, classifying unbiased larvae was less accurate, yet a single 4x trial accurately represented 10/18 (55.56%) of individuals. Altogether, the q4X testing strategy confirms our earlier findings and demonstrates the veracity of our recording strategies to detect specific turning types.

As the q4X assay allowed for a rigorous confirmation of turning type, we next wanted to determine whether left, right, or unbiased turning types exhibited unique path trajectory characteristics. A PI was calculated from all 16 trials in the q4X assay for each individual and categorized as left, unbiased, or right type. For each turning type, we examined fractal dimension (F.D.) and displacement (displ) measure of localized movement density (Tremblay et al., 2007; Horstick et al., 2017). Comparison across all three motor types yielded no differences in the tested motor parameters (main effect due to turn type 2-way ANOVA displ: *F(2,222) = 0*.*42*, p=0.66; F.D: *F(2,222) = 2*.*12*, p=0.12), yet the expected changes in behavior following light extinction were observed (main effect due to illumination 2-way ANOVA displ: *F(1,222) = 604*, p<0.0001 ; F.D: *F(1,222) = 643*, p<0.0001) (Figure 2F-G). Interestingly, upon closer inspection, we did notice a small yet significant change in F.D. between left and right turning groups during dark trials (left 1.240 ± 0.012; right 1.200 ± 0.014: *t(222)=2*.*974*, adjusted p=0.0489, effect size d=0.63). This effect was specific, and not observed during baseline (left F.D. 1.021 ± 0.003; right F.D. 1.021 ± 0.004: *t(222)=0*.*059*, adjusted p>0.9999) or for displacement. These results show that the difference of left and right turning type also generate mild changes to search pattern behavior, yet not motor trajectories during baseline movement.

### Development of inter-individual variation is sensitive to specific environmental factors

Many instances of motor and behavioral biases show limited heritability (COLLINS, 1969; Buchanan et al., 2015; Linneweber et al., 2020). This observation suggests that inter-individual variation is, at least in part, modulated through individual experience with environmental factors. Indeed, previous studies show that social experience and environmental enrichment modify inter-individual variation of specific behaviors (Freund et al., 2015; Akhund-Zade et al., 2019; Versace et al., 2020; Zocher et al., 2020). As larval zebrafish turning bias is not heritable (Horstick et al., 2020), we reasoned that the environment might contribute to overall inter-individual variation or the generation of specific turning types. To test this hypothesis, we first established that turn bias appears at 4 dpf (Supplementary Figure 3A-B). Therefore, larvae were exposed to changes in the environment from 1 through either 4 or 7 dpf, dependent on the tested factor. The four parameters we screened were social experience, environmental enrichment, temperature, and salinity (Figure 3A). Social interaction and environmental enrichment were selected because each has been shown to modulate inter-individual variation (Akhund-Zade et al., 2019; Versace et al., 2020). For social interaction, larvae are raised in isolation or groups. For enrichment, we generated two environments: 1) an enriched environment where a petri dish was fitted with internal surfaces, diverse color, hiding spots, water surface cover, and dynamic substrate pattern, and 2) an empty petri dish with a uniform white bottom as a control. In addition, we also tested the impact of etiologically relevant temperature (24 or 32°C) and salinity (0.5-5 parts per thousand (ppt)) variations during early development compared to standard rearing parameters (Engeszer et al., 2007; Sundin et al., 2019). Thus, our parameters test factors that generated inter-individual variation in other models and abiotic environmental fluctuations that larvae could encounter in a native habitat.

**Figure 3.**
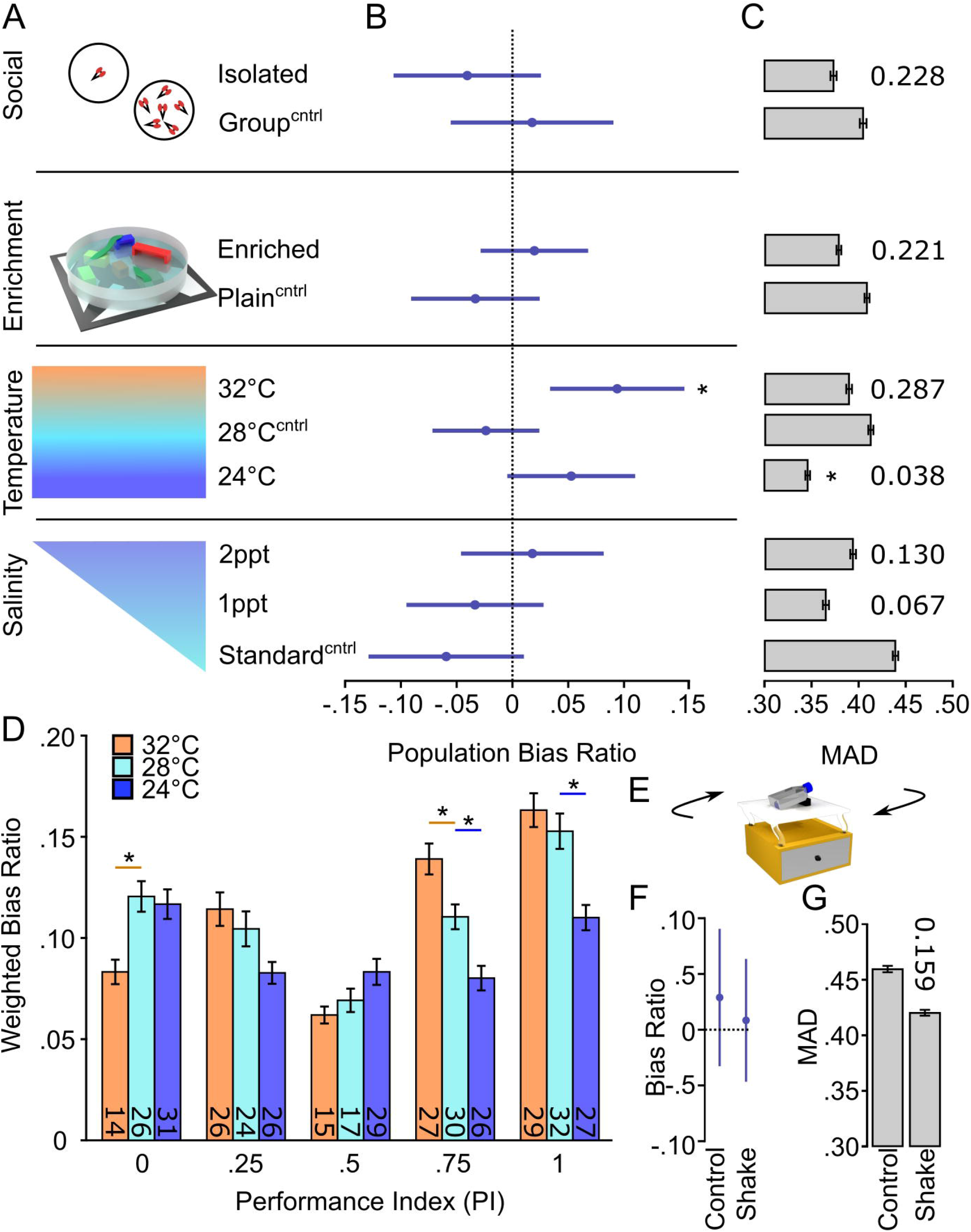
Temperature selectively changes inter-individual variation in turn bias. **(A)** Schematic of environmental manipulations (Social Isolated N= 87, Social Group N=87; Enriched N=172, Plain N=134; Temp. 32°C N=112, Temp. 28°C N=136, Temp. 24°C N=144; Standard salinity N=102, 1ppt N=107, 2ppt N= 107). Each condition has an independent control denoted by superscript Cntrl. **(B)** Average bias ratios across the entire tested population per condition. Statistical comparison performed to 0, denoting no population level bias. **(C)** MAD for dark responses, error bars are 95% confidence levels generated from 1000 bootstraps. Individual p-values shown above bars, calculated by comparing experimental groups to controls. **(D)** Average bias ratio per PI weighted by the percent of individuals. N’s indicated within bars. **(E)** Representative diagram of the setup used to shake larvae during early development. Average population bias ratio **(F)** and MAD **(G)** between shake experiments. Number above bar represents p-value compared to the control group. Asterisk p <0.05.

To determine if any of the tested parameters altered turning type development or magnitude of inter-individual variation, we looked at the average population bias ratio and MAD, respectively (Figure 3B-C) (Supplementary Figure 3C). Interestingly, the elevated temperature during early development caused a significant population shift from random (high temp 0.094 ± 0.044: one-sample *t*-test against 0, *t(112)* = *2*.*157*, p=0.033), implicating a population-level rightward bias, whereas no significant changes were observed in other temperature conditions or any other tested environmental parameter (Figure 3B). Conversely, the magnitude of turn bias variation during light-search was only reduced by low-temperature rearing, yet unaffected by other testing conditions (Figure 3C). To confirm the observed temperature-dependent changes, we examined the bias ratio per PI, weighted for the number of individuals per PI group. We observed that temperature imposed a significant effect on turn bias persistence (main effect of temperature 2-way ANOVA *F(2,364) = 9*.*275*, p=0.0001) (Figure 3D). Indeed, the tested high temperature resulted in a significant depression of leftward turning (within PI group comparison *t(364) = 3*.*031*, adjusted p=0.0078; red line) and increase in rightward turning (0.75 PI *t(364) = 2*.*904*, adjusted p=0.012; red line). Conversely, low temperature depressed turn bias performance in the population (Figure 3D, blue lines). These results suggest a specific temperature-mediated change. However, in larval zebrafish, fluctuating temperature is a stressor, and elevation of stress signaling has been shown to attenuate visual bias in chickens (Rogers and Deng, 2005; Long et al., 2012; Haesemeyer et al., 2018). Therefore, we tested the effect of shaking on turn bias which is a potent stressor for larval zebrafish (Eto et al., 2014; Castillo-Ramírez et al., 2019; Apaydin et al., 2020). Sustained shaking during early development resulted in no population or turn bias magnitude changes (Figure 3E-F panels). Moreover, external temperature impacts the rate of zebrafish development, and based on previous studies, our conditions would lead to an estimated ±13 hour shift in development (Kimmel et al., 1995). We show that our temperature assay results in a change in hatching, a developmental marker, yet no gross changes in morphology or survival (Supplementary Figure 3D-F). These data illustrate that etiologically relevant temperature fluctuations differentially and specifically affect inter-individual turn bias variation.

### Elevated temperature impacts rostral PT specification

A basic circuit involving the rostral posterior tuberculum (PT) and dorsal habenula (dHb) neurons has previously been described for zebrafish turn bias (Horstick et al., 2020). However, in wildtype larvae, no hemispheric differences in these neurons were found to account for left or right turning preference (Horstick et al., 2020). Because we found that elevated temperature disrupted left and right turning balance, we next wanted to determine if elevated temperature caused changes in neurons necessary for turn bias. We reasoned our environmental variables could alter neuronal development, as bias maintaining PT neurons are present as early as 2 dpf (Horstick et al., 2020), and dHb differentiation begins on 1 dpf (Gamse et al., 2003; Amo et al., 2010). First, we wanted to identify if a specific period during early development was sensitive to increased temperature. We found that elevated temperature during either 31-55 hpf or 55-79 hpf intervals did not recapitulate the population shift observed during the 1-4 dpf exposure (Supplementary Figure 4); therefore, we selected the full testing duration for further investigation. To visualize key dHb and PT neurons, we used the enhancer trap line *y279:Gal4*, which labels both populations of neurons (Horstick et al., 2020) (Figure 4A). In zebrafish, the left dHb is considerably larger than the right dHb (Gamse et al., 2003; Roussigné et al., 2009). We found that elevated temperature did not alter the habenula, and typical left/right asymmetry was observed (2-way ANOVA: interaction between temperature and hemisphere *F(1,56) = 0*.*549*, p=0.46) ; effect of hemisphere *F(1,56) = 91*.*70*, p<0.0001) (Figure 4B,E). No hemispheric differences (main effect of hemisphere 2-way ANOVA *F(1,56) = 0*.*427*, p=0.52) were observed in the number of y279 positive PT neurons (Figure 4C). Therefore, we combined PT measures from both hemispheres. Interestingly, from these combined pools, y279 positive neuron area in the PT was reduced after exposure to elevated etiological temperature during early development (high temperature 255.65 ± 17.73; normal temperature 348.50 ± 20.84: 2-tail *t*-test *t(58) = 3*.*385*, p=0.0013) (Figure 4D,F), establishing a potential neuronal basis for how high temperature during development modifies turn bias inter-individual variation.

**Figure 4.**
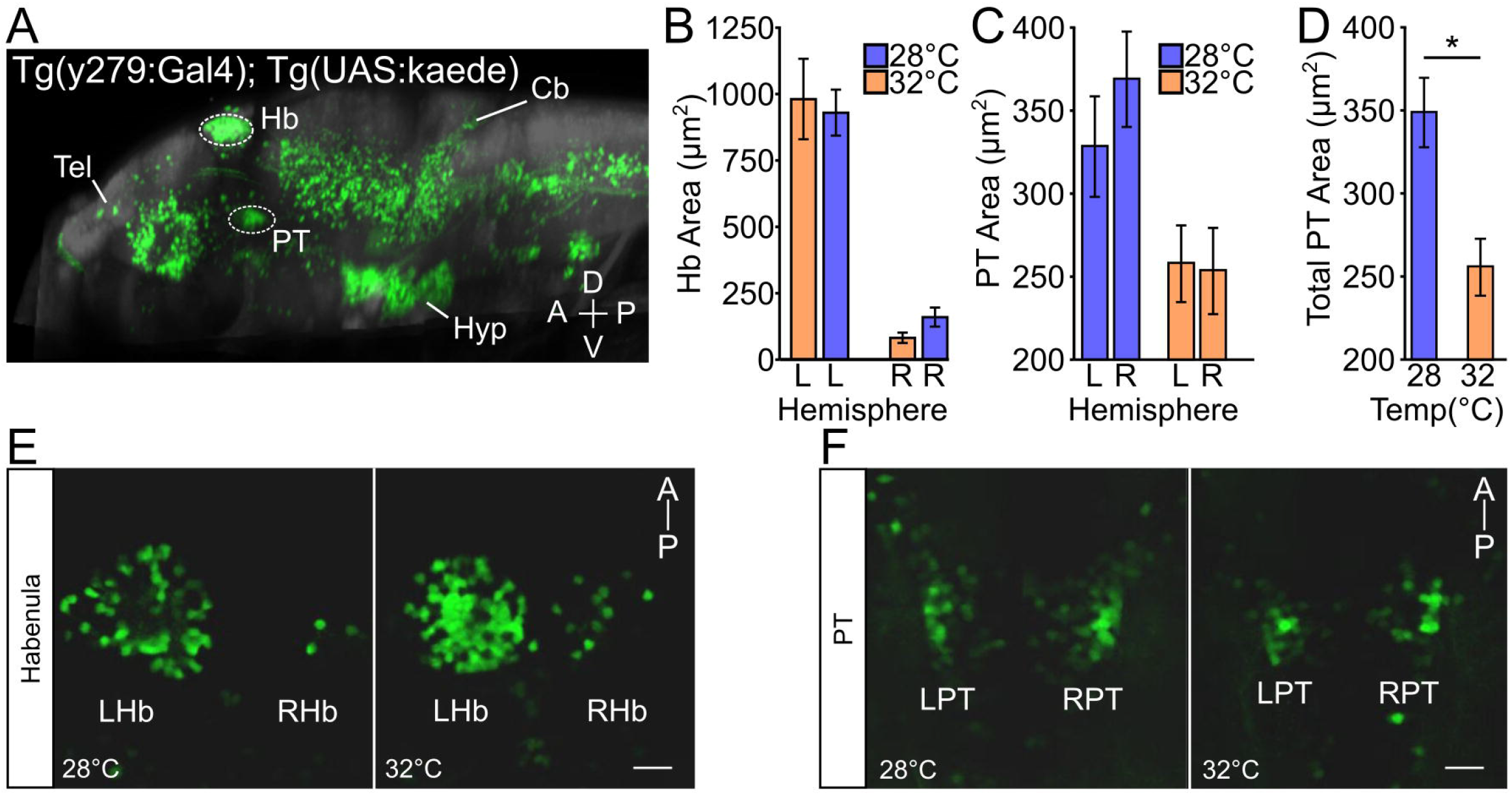
Temperature impacts y279 specified expression in the PT. **(A)** Sagittal view of larval zebrafish brain showing expression of enhancer trap *Tg(y279:Gal4)* obtained from the zebrafish brain browser atlas. Circled regions highlight the habenula (Hb) and rostral posterior tuberculum (PT) and white lines show the telencephalon (Tel), hypothalamus (Hyp), and cerebellum (Cb). **(B-F)** Effect of elevated temperature during early development on the expression of y279 in the habenula and PT. (**B)** Expression of y279 in the left and right hemisphere Hb nuclei (28°C purple, N=16; 32°C orange, N=14). (**C)** Hemispheric area of y279 positive PT neurons (28°C purple, N=16; 32°C orange, N=14). (**D)** Combined left and right hemisphere PT areas (28°C purple, N=16; 32°C orange, N=14). **(E-F)** Representative images for y279 positive Hb (left habenula, LHb; right habenula, RHb) **(E)** and PT (left PT, LPT; right PT, RPT) **(F)** neurons for larvae raised at 28°C or 32°C. Scale bar 20 µm. Asterisk p<0.05.

### Motor individuality is sensitive to gene signaling associated with neuronal proliferation

Studies from *C. elegans* (Bertrand et al., 2011) and *Drosophila* (Linneweber et al., 2020) demonstrate that Notch signaling can generate functional asymmetries in the brain that drive unique individual behavioral responses. Established zebrafish mutant lines *mindbomb (mib)* and *mosaic eyes (moe)*, E3 ubiquitin ligase and Epb41l5 adapter respectively, do not directly disrupt the Notch cascade, yet impair Notch signaling (Itoh et al., 2003; Ohata et al., 2011; Matsuda et al., 2016). Indeed, haploinsufficiency in these lines abrogates zebrafish turn bias, suggesting sensitivity to the levels of Notch signaling (Horstick et al., 2020). One of the canonical roles of Notch during early development is the regulation of neuronal proliferation (Appel et al., 2001; Mizutani et al., 2007; Yoon et al., 2008). Therefore, we next aimed to elucidate if turn bias is 1) sensitive to direct Notch antagonism in a dose-dependent manner and 2) if partial Notch inhibition impairs neuronal proliferation.

To disrupt Notch signaling, we used the specific γ-secretase inhibitor LY411575, which blocks the activation of the Notch signaling cascade (Geling et al., 2002; Fauq et al., 2007). Previous reports show that treatment with micromolar concentrations of LY411575 starting at mid-gastrulation results in a near-total loss of Notch signaling, which largely recapitulates the *mindbomb* mutant (Jacobs and Huang, 2019; Sharma et al., 2019). Therefore, we used 10 µM as a maximum dose and positive control for inhibitor efficacy across trials. To identify a level of Notch inhibition that could impair turning bias, we LY411575-treated larvae from mid-gastrulation to 4 dpf over 7 concentrations ranging from 50nM to 10 µM and scored phenotypes at 3 dpf (Figure 5A). Developmental exposure of LY411575 up to 100nM left most larvae morphologically normal, which we used as a maximum dose to test the impact on turn bias. Notch inhibition resulted in a significant change in TTA following the loss of illumination (1-way ANOVA *F(2,152) = 4*.*614*, p=0.011), causing an increase in overall turning at 100nM inhibitor treatment compared to controls (vehicle 1175.95 ± 53.34, 100nM 1411.39 ± 66.50: *t(152) = 2*.*786*, adjusted p=0.018) (Supplementary Figure 5). Whereas turn bias performance was reduced by Notch inhibition (main effect due to treatment 2-way ANOVA *F(2,144) = 8*.*995*, p=0.0002), with 100nM inhibitor nullifying bias ratio strength differences due to PI, which was not observed at lower inhibitor concentrations (Figure 5B-C). In addition, 100nM but not 50nM treatment reduced overall inter-individual turn bias variation in the population (Figure 5D). This data suggests that a critical threshold of Notch signaling is required for generating variation in turn bias and overall performance, which is lower than levels necessary for normal gross morphological development. To confirm that LY411575 exposure impaired Notch signaling, we examined *her12* expression, a downstream target of the Notch signaling cascade that is robustly expressed in the spinal cord, providing an unambiguous region to quantify expression changes (Jacobs and Huang, 2019). Exposure to micromolar inhibitor concentrations resulted in a near-total absence of *her12* expression, consistent with previous reports (Jacobs and Huang, 2019). The *her12* expression was, however, observed in the spinal cord of the 100nM group at an intensity indistinguishable from controls (Figure 5E, G).

**Figure 5.**
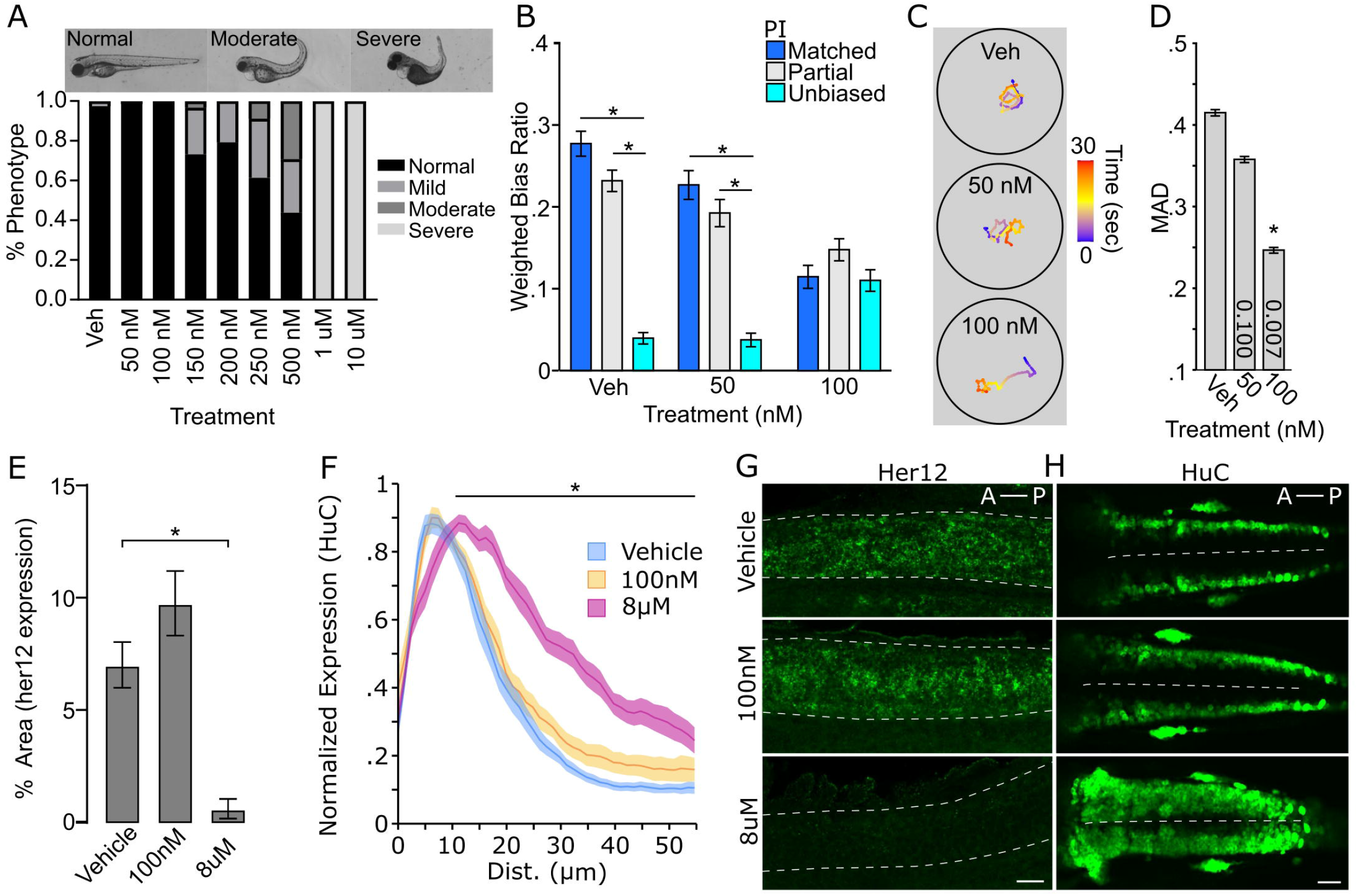
Turn bias is sensitive to levels of Notch signaling. **(A)** Phenotypic counts following Notch inhibitor treatment (Vehicle, N=125; 50nM, N=88; 100nM, N=114; 150nM, N=123; 200nM, N=120; 250nM, N=101; 500nM, N=48; 1µM, N=43; 10µM, N=100). **(B)** Weighted absolute bias ratio averages (matched PI=0,1 blue bar; Partial match PI= 0.25, 0.75 grey bar; unbiased PI=0.50 cyan bar. **(C)** Illustrative traces for treatment groups. Scale bar color represents time in seconds. **(D)** Effect of Notch inhibition on MAD. p-values shown in bar, 1-way comparison of treatment groups to control. **(E)** Area of her12 expression in the spinal cord following LY411575 treatment (Vehicle, N=12; 100nM, N=13; 8uM, N=11). **(F)** Normalized distribution of HuC/D positive neurons following notch inhibition (Vehicle: Blue, N=18; 100nM: Yellow, N=13; 8µM: Magenta, N=13). X-axis distance spans half the spinal cord (0 micron = lateral spinal cord; 55 micron = spinal cord midline). Comparison shown is between vehicle and 8µM along the whole length of black bar between matched positions. Ribbons ±SEM. **(G)** Representative images of her12 expression in 27 hpf embryos. Lateral view of spinal cord (dotted outline). Scale bar 20µm. **(H)** Representative HuC labeling in 24 hpf embryos showing dorsal view. Dotted line denotes spinal cord midline. Scale Bar 40µm. Asterisk p<0.05.

A canonical and conserved role for Notch during early development is regulating neuronal proliferation and maintaining progenitor pools, and the loss of Notch leads to increased proliferation (Appel et al., 2001; Cheng et al., 2004; Sharma et al., 2019). Therefore, we next wanted to determine whether the level of Notch inhibition that impairs turn bias individuality also disrupts proliferation. During zebrafish embryonic development, proliferative neurons are readily visualized in the anterior hindbrain using Elav (HuC/D) protein expression as a marker (Kim et al., 1996; Sharma et al., 2019). These proliferative neuron pools expand following high levels of Notch inhibition or in the *mindbomb* mutant background (Itoh et al., 2003; Sharma et al., 2019). Similar to our observation with *her12*, partial pharmacological Notch inhibition (100nM drug) induced no change in actively proliferating neurons, yet positive controls (8µM) displayed robust expansion of proliferating neurons (Figure 5F,H).

Notch signaling is ubiquitous in the larval zebrafish nervous system (Tallafuss et al., 2009; Banote et al., 2016; Kumar et al., 2017), and pharmacological inhibition is not specific. Consequentially, we next aimed to determine whether proliferative pathways in restricted areas of the brain may also contribute to turn bias. Genomic screen homeobox transcription factors (Gsx1 and 2, formerly Gsh1 and 2) are affecters of the Notch signaling pathway in mouse, and Gsx2 maintains neural progenitor pools in the developing telencephalon (Wang et al., 2009; Pei et al., 2011; Roychoudhury et al., 2020). In larval zebrafish, *gsx2* is predominantly expressed in the pallium, preoptic area, hypothalamus, and hindbrain, with an established putative null TALEN deletion mutant line (Coltogirone et al., 2021). As *gsx2* mutants show no gross morphological abnormalities during larval stages, we used these lines to test turn bias. Heterozygous and mutant *gsx2* larvae displayed reduced inter-individual variation and a shift toward less persistent turn bias (Figure 6A). The loss of persistent same-direction turning was similarly observed using match index (MI), an analogous metric (Figure 6B). Yet, TTA during light-search was not significantly changed across genotypes (1-way ANOVA *F(2,187) = 2*.*730*, p=0.068), suggesting the loss of same-direction turning is not due to reduced light-driven behavior (Figure 6C). Thus, our analysis implies that broad and local haploinsufficient changes in Notch signaling and Gsx2 contribute to inter-individual variation in turn bias behavior.

**Figure 6.**
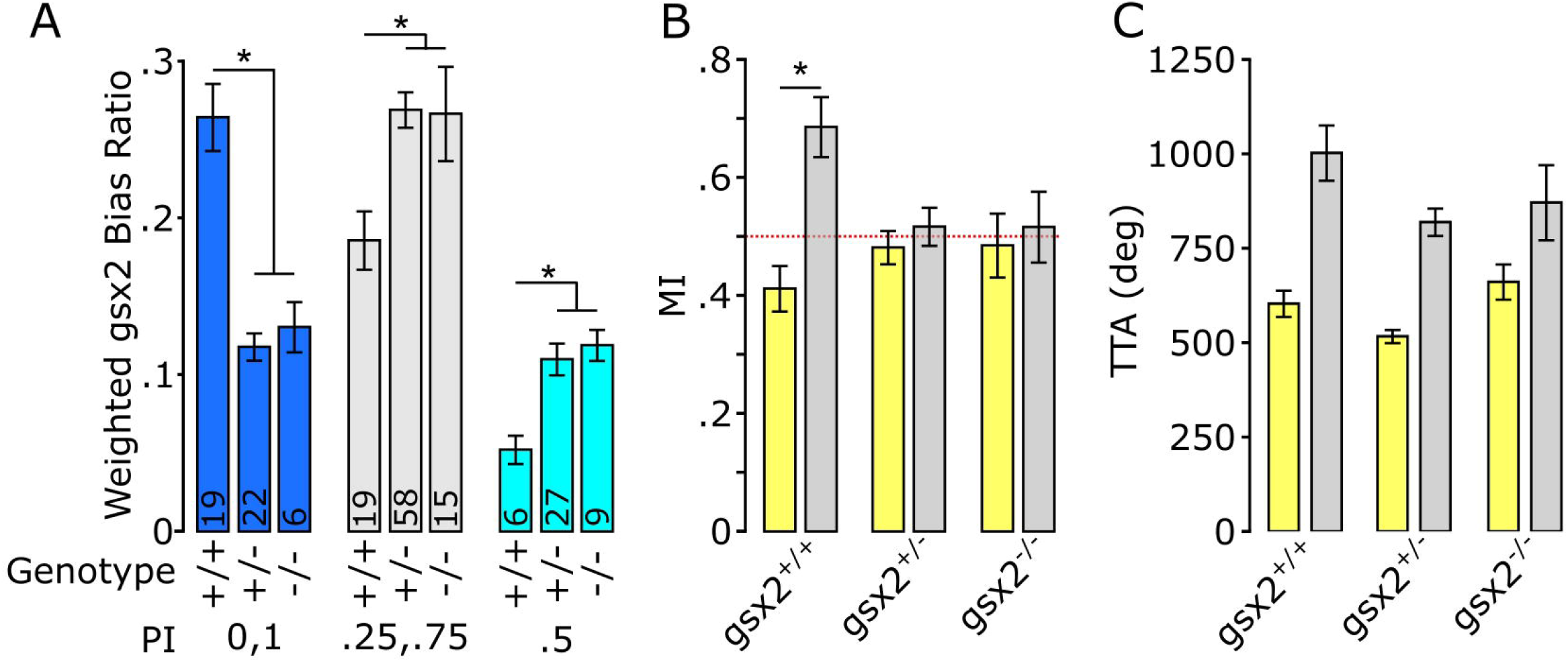
Loss of *gsx2* reduces inter-individual variation. **(A)** Effect of gsx2 genotype on weighted average bias ratio per performance groups (Matched: PI=0,1; Partial PI=0.25,0.75; Unbiased PI=0.5). Numbers on bars indicate N. Weighing was based on the percent of larvae within each PI per genotype. **(B)** MI shows that only gsx2 +/+ larvae maintain persistent same-direction turning following the loss of illumination. Dotted line at 0.5 indicates random movement **(C)** TTA between genotypes is not affected (gsx2 +/+, N=45; gsx2 +/-, N=113; gsx2 -/-, N=32). Baseline (yellow) and dark (grey) responses in B-C. Asterisk p<0.05

## Discussion

Here we reveal that during light-search initiated by the loss of illumination, larval zebrafish exhibit significant inter-individual variation in turn bias, a handed-like behavior. Based on our newly developed assays, we were further able to show mild changes in search behavior correlated with left and right turning types. However, the impact of turning on search motor patterns was specific, as we found no evidence of individual motor types changes during baseline illumination, consistent with previous studies (Horstick et al., 2020). We demonstrated a turn bias spectrum across the population which shows the previously described left/right turning types (Horstick et al., 2020). In addition, our analysis revealed a consistently unbiased turning type, supported by multiple independent recording strategies (4X, 8X, and q4x). Furthermore, we show that temperature changes during early development result in sustained changes in inter-individual variation. Finally, we tested how signaling pathways associated with neuronal proliferation affected turn bias development, using either pharmacological inhibition of Notch signaling or a presumable null Gsx2 mutant. Notch and Gsx2 represent canonical broad and regional regulators of proliferation, respectively. Interestingly, turn bias attenuation is observed with partial Notch inhibition and in *gsx2* heterozygotes, suggesting dose-dependent sensitivity. Despite a well-established role for Notch in cell proliferation, the partial inhibition that selectively impairs turn bias did not result in observable changes, at least early in development (see Figure 5). Our findings confirm that three turning types can be categorically defined, are modulated by specific etiological relevant environmental cues, and are sensitive to internal proliferative associated signaling pathways. One potential caveat is that zebrafish strains are not isogenic and maintain some genetic heterogeneity (Butler et al., 2015), potentially contributing to inter-individual differences. Nevertheless, our work develops larval zebrafish as a powerful model to identify mechanisms generating inter-individual variation in vertebrates.

### Determination of bias

Our findings suggest a ‘hemispheric noise’ model where turn bias and inter-individual variation is modulated by conflicting signals in turn bias driving neurons between the brain hemispheres (Figure 7). We elucidated that change in bias ratios strength distinguishes unbiased versus biased larvae. Moreover, we establish that this change is not a result of a loss of photo-responsiveness in unbiased individuals (total turning, see Supplementary Figure 1); rather a failure to navigate in a single direction during light-search consistently. This observation supports the conclusion that unbiased individuals are not a subset with impaired photo-responsiveness, but a distinct behavioral motor profile during search behavior. Supporting this model, when we quantify the strength of individual trials, the bias ratios exhibit a step-wise decrease, i.e., PI 1<0.75<0.5, suggesting accumulating inter-hemispheric noise degenerates overall individual bias persistence. Corroborating this model, previous studies showing that unilateral ablation of rostral PT neurons, which are required for turn bias in larval zebrafish, increases turning strength in the direction ipsilateral to the intact neurons, indicating ablation removes conflicting input from the contralateral hemisphere (Horstick et al., 2020). In pigeons, a classic model for hemispheric specialization and individual variation (Güntürkün et al., 1998; Freund et al., 2016), increased conflict between hemispheres exacerbates visual task latency (Manns and Römling, 2012). Therefore, variable balance in hemispheric signaling may be a conserved mechanism in generating inter-individual variation (Chen-Bee and Frostig, 1996; Linneweber et al., 2020). Inter-hemispheric communication is vital for the function of the visual system (Bui Quoc et al., 2012; Chaumillon et al., 2018), including photo-driven behavior in larval zebrafish (Gebhardt et al., 2019). The counter hypothesis is a ‘switching model’ where unbiased larvae display vigorous directional turning in randomly selected directions over sequential trials. This model is consistent with a ‘winner take all’ circuit function (Fernandes et al., 2021). Indeed, within the primary visual processing center in zebrafish, the optic tectum, neurons operate in a winner take all style during visually guided behavior (Fernandes et al., 2021). However, turn bias is driven by the loss of visual cues that activate rostral PT neurons, which do not project to the tectum (Horstick et al., 2020), implicating that even though turn bias is evoked by visual input, the mechanism is likely independent of a tectal winner take all mechanism. Despite the neurons maintaining zebrafish turn bias being identified, the underlying mechanism imposing a specific turning type remains unknown (Horstick et al., 2020). Our analysis suggests a model of competitive inter-hemispheric communication modulating the magnitude of inter-individual turn bias variation that is further adjusted by fluctuating and specific variables in the internal and external environment.

**Figure 7.**
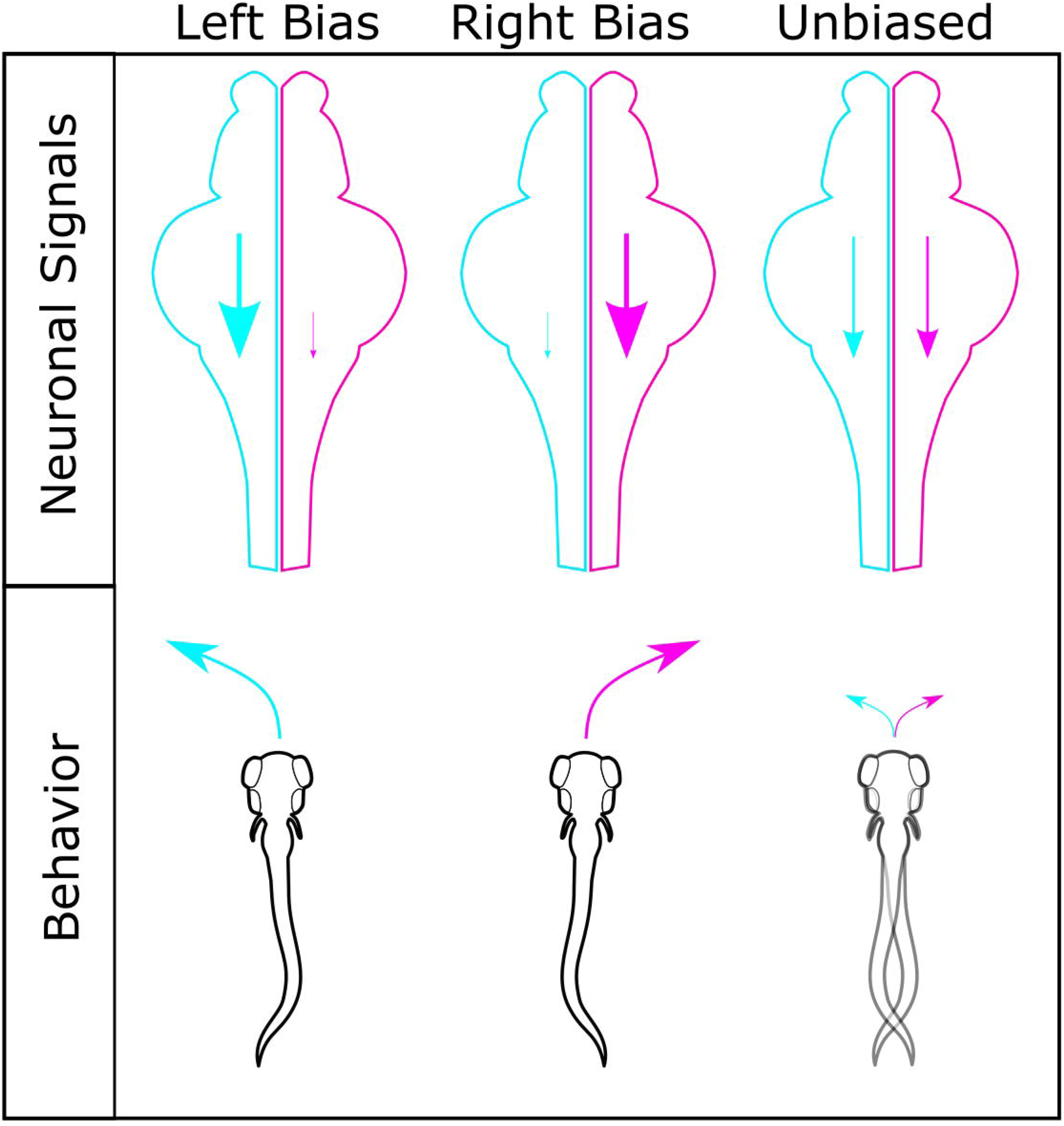
Model for generating different turning types. Interhemispheric differences in turn bias driving motor signals are a potential mechanism for establishing turning types. Left (cyan) and right (magenta) hemispheres shown for left, right, and unbiased motor types, with corresponding motor drive shown by scale of descending arrow. For individuals with robust left or right turning types, a strong ipsilateral turn bias signal persists in a single hemisphere, with limited conflicting input from the contralateral hemisphere. In individuals with similar turn bias drive from both hemispheres, no single turn direction persists due to conflict between hemispheres, resulting in lower bias ratios and unbiased turning types.

### Regulation of individuality

The mechanisms driving unique individual behavioral responses based on sex or sensory context are well described (Asahina et al., 2014; Lewis et al., 2015; Yapici et al., 2016; Marquart et al., 2019; Ishii et al., 2020; Nelson et al., 2020). However, the basis of individuality that is observed in many species to consistent stimuli is still poorly understood, especially in vertebrates. One of our goals was to determine what internal and external environment elements may modulate turn bias variation. To test external environmental factors, we selected environmental enrichment, social experience, temperature, and salinity. One hypothesis for how factors like enrichment or social experience influence inter-individual variation is through micro-environmental interactions that create unique individual experiences (Kain et al., 2015). However, our data indicate that these interactions do not influence turn bias variation in zebrafish. One possible explanation is that during early development, 1-3 days post-fertilization, larvae are primarily inactive and only begin actively exploring around 4 days post fertilization (Colwill and Creton, 2011; Lambert et al., 2012). Conversely, responsiveness to conspecifics is not observed until 3 weeks (Dreosti et al., 2015; Larsch and Baier, 2018). As bias is established by 4 dpf, the underlying mechanisms may no longer be malleable to environmental experiences beyond this developmental interval.

We also tested temperature and salinity, emphasizing etiological ranges that zebrafish could experience in their native environments (Engeszer et al., 2007; Sundin et al., 2019). Salinity and temperature are critical environmental determinants and have been shown to drive evolutionary changes in stickleback populations (Gibbons et al., 2017). However, we found that only raising larvae at varying temperatures resulted in modifications to inter-individual variation. We show that temperature-dependent effects are not a generic thermal stress response. Etiological increases in temperature have been shown to attenuate turn bias in adult reef fish, implicating a potentially broader thermal sensitivity in bias establishing mechanisms (Domenici et al., 2014). Our analysis establishes that early developmental exposure to etiological temperature fluctuation results in sustained and specific turn bias changes.

Intriguingly, the specification of habenular hemispheric asymmetry is sensitive to the rate of development (Aizawa et al., 2007), and developmental rate is temperature sensitive (Kimmel et al., 1995). This observation could provide a potential mechanism for thermal generated changes in turn bias. However, our testing conditions produced no gross change in the habenula nuclei morphology. This observation, however, does not exclude functional or subcellular changes. Surprisingly, we observe a bilateral reduction in *Tg(y279)* positive PT neurons, which are essential for maintaining turn bias, in the elevated temperature experiments. A primary function of the PT is to integrate diverse sensory inputs (Striedter, 1991; Derjean et al., 2010; Yaeger et al., 2014). However, thermosensitivity of the PT neurons has not been previously described, and we believe this is a novel observation. Future studies identifying the genetic basis of the *Tg(y279)* enhancer trap, which is currently unknown, will be instrumental in elucidating how temperature changes PT neuron specification and inter-individual variation. The specific abrogation of leftward turning types in increased temperature conditions provides a powerful model to interrogate underlying neural changes in a vertebrate brain associated with individual behavioral patterns.

Last, we wanted to identify molecular signaling pathways regulating turn bias. Biased turning in larvae is largely lost in heterozygotes of mutant lines associated with Notch signaling, yet the impact of direct Notch inhibition was unexplored (Horstick et al., 2020). In *Drosophila* and *C. elegans*, Notch signaling is essential for establishing individual visual navigational strategies and asymmetric chemosensory neuron identities, respectively (Bertrand et al., 2011; Linneweber et al., 2020). Thus, work from several species implicates Notch as a driver of variation at behavioral and neuronal levels. Indeed, we show that partial Notch disruption, using a specific pharmacological inhibitor, disrupts biased turning in larval zebrafish, yet not the ability to respond to illumination changes, establishing a direct role of Notch signaling for turn bias independent of morphological development. Despite the established role of Notch in neural proliferation, we found no significant change in proliferative neurons or *her12* expression at the dosages used for behavioral studies. Since Notch signaling is essential for diverse cellular functions, and the precise Notch mediated signaling mechanisms are highly sensitive to the strength of Notch signaling (De Smedt et al., 2005; Shen et al., 2021), the low inhibitor concentrations used may be sub-threshold for disturbing the spatial-temporal patterns of *her12* and HuC/D tested here. In addition, the downstream effects of Notch are dependent on the cellular micro-environments, such as the co-expression of Notch receptors, ligands, and auxiliary proteins (Demehri et al., 2009; Bertrand et al., 2011; LaFoya et al., 2016). Therefore, the levels of Notch reduction that impair turn bias, but not morphology, may not be sufficient to alter Notch associated mechanisms impacting proliferation. Nevertheless, subtle changes in Notch could lead to changes in cellular micro-environments, thereby altering downstream signaling cascades, and ultimately impacting turn bias maintaining neurons. Notch haploinsufficiency is known to generate a myriad of defects and disease states, including vasculature defects, seizure, autism, and brain malformations, demonstrating that reduced Notch signaling can disrupt biological functions (Krebs et al., 2004; Connor et al., 2016; Fischer-Zirnsak et al., 2019; Blackwood et al., 2020). However, the pharmacological inhibition used in our current study is not regionally specific. Therefore, we also tested an established zebrafish *gsx2* mutant line, and *gsx2* is predominately expressed in subsets of hypothalamic, preoptic area, pallium, and hindbrain neurons (Coltogirone et al., 2021). The reduction in turn bias in *gsx2* heterozygotes and mutants suggests that turn bias variation is sensitive to local changes in brain regions where *gsx2* is expressed, independent of the previously described rostral PT and habenula (Horstick et al., 2020). As the levels of Notch that reduce turn bias do not impact proliferation, it seems possible that Notch and Gsx2 modulate turn bias by independent mechanisms. Our current analysis identifies two conserved molecular signaling and transcriptional control mechanisms, Notch and Gsx2, and novel neuroanatomical substrates as important for generating variation in turn bias.

### Function of turn bias and inter-individual variation

Behavioral variation is observed in diverse species and behavioral modalities (Byrne et al., 2004; Elnitsky and Claussen, 2006; Cauchard et al., 2013; Horváth et al., 2020). In zebrafish, even complex neuromodulatory processes such as startle habituation display inter-individual variation with distinguishable ‘habituation types’ (Pantoja et al., 2016, 2020). Yet, the general question remains, “why do specific behavioral modalities manifest inter-individual differences?” Considering a simple form of inter-individual variation, such as turn bias, may offer insights to these questions. Zebrafish are active hunters during larval stages and predatory success depends on visual input, thus establishing a potent drive to remain in illuminated areas (Gahtan et al., 2005; Filosa et al., 2016; Muto et al., 2017). Following the loss of light and of overt navigation cues, larvae initiate a local light-search, where individual turn bias is triggered, causing looping trajectories (Horstick et al., 2017). Looping search trajectories are observed in various species in the absence of clear navigational cues, suggesting an efficient systematic strategy (Collins et al., 1994; Conradt et al., 2000; Zadicario et al., 2005). However, even seemingly optimal behaviors may not be advantageous in all contexts (Simons, 2011). Variation in turning types may ensure individuals across the population possess strategies to mitigate various environmental challenges, a form of bet-hedging (Simons, 2011; Kain et al., 2015). Similarly, behavioral variation adds unpredictability to a population. Predictable behavioral patterns can be exploited by predators (Catania, 2009, 2010). For aquatic species, this may be advantageous as some heron species, a predator of small fish, use a canopy hunting strategy, covering the water surface with their wings and blocking light (Kushlan, 1976). Prey populations with unpredictable responses would potentially provide a more challenging target (Humphries and Driver, 1970). Even though larval fish may not be the intended target of heron canopy hunting, larval behavioral patterns may persist over their lifespan. Indeed, adult zebrafish display a persistent turn direction preference (Fontana et al., 2019), although the correlation to larval turn bias is currently unexplored. Ultimately, the etiological purpose for turn bias variation is most likely a combination of multiple explanations, including bet-hedging, generating unpredictability, and genetically encoded sources of variation.

## Author Contributions

E.J.H conceived the experiments. E.J.H and J.H wrote the manuscript and analyzed data. S.L wrote custom R scripts for data analysis. S.B provided *gsx2* lines and associated analysis. J.H, M.W, J.S, H.C, H.P, L.B, C.S performed experiments.

## Acknowledgements

We thank Alexandra Schmidt and Rebecca Coltogirone from the Bergeron lab for help with imaging and fluorescent in-situs and Erik Duboué and Andrew Dacks for helpful comments on the manuscript.

## Data Availability

All datasets and custom R scripts generated in this study will be available upon request.

## References

Aizawa, H., Goto, M., Sato, T., and Okamoto, H. (2007). Temporally Regulated Asymmetric Neurogenesis Causes Left-Right Difference in the Zebrafish Habenular Structures. Developmental Cell 12, 87–98. doi:10.1016/j.devcel.2006.10.004.

Akhund-Zade, J., Ho, S., O’Leary, C., and de Bivort, B. (2019). The effect of environmental enrichment on behavioral variability depends on genotype, behavior, and type of enrichment. Journal of Experimental Biology 222. doi:10.1242/jeb.202234.

Amo, R., Aizawa, H., Takahoko, M., Kobayashi, M., Takahashi, R., Aoki, T., et al. (2010). Identification of the Zebrafish Ventral Habenula As a Homolog of the Mammalian Lateral Habenula. J. Neurosci. 30, 1566–1574. doi:10.1523/JNEUROSCI.3690-09.2010.

Andrew, R. J., Dharmaretnam, M., Gyori, B., Miklósi, A., Watkins, J. A. S., and Sovrano, V. A. (2009). Precise endogenous control of involvement of right and left visual structures in assessment by zebrafish. Behav. Brain Res. 196, 99–105. doi:10.1016/j.bbr.2008.07.034.

Apaydin, D. C., Jaramillo, P. A. M., Corradi, L., Cosco, F., Rathjen, F. G., Kammertoens, T., et al. (2020). Early-Life Stress Regulates Cardiac Development through an IL-4-Glucocorticoid Signaling Balance. Cell Reports 33, 108404. doi:10.1016/j.celrep.2020.108404.

Appel, B., Givan, L. A., and Eisen, J. S. (2001). Delta-Notch signaling and lateral inhibition in zebrafish spinal cord development. BMC Developmental Biology 1, 13. doi:10.1186/1471-213X-1-13.

Asahina, K., Watanabe, K., Duistermars, B. J., Hoopfer, E., González, C. R., Eyjólfsdóttir, E. A., et al. (2014). Tachykinin-Expressing Neurons Control Male-Specific Aggressive Arousal in Drosophila. Cell 156, 221–235. doi:10.1016/j.cell.2013.11.045.

Ayroles, J. F., Buchanan, S. M., O’Leary, C., Skutt-Kakaria, K., Grenier, J. K., Clark, A. G., et al. (2015). Behavioral idiosyncrasy reveals genetic control of phenotypic variability. Proc Natl Acad Sci USA 112, 6706–6711. doi:10.1073/pnas.1503830112.

Banote, R. K., Edling, M., Eliassen, F., Kettunen, P., Zetterberg, H., and Abramsson, A. (2016). β-Amyloid precursor protein-b is essential for Mauthner cell development in the zebrafish in a Notch-dependent manner. Developmental Biology 413, 26–38. doi:10.1016/j.ydbio.2016.03.012.

Bell, W. J., Cathy, T., Roggero, R. J., Kipp, L. R., and Tobin, T. R. (1985). Sucrose-stimulated searching behaviour of Drosophila melanogaster in a uniform habitat: modulation by period of deprivation. Animal Behaviour 33, 436–448. doi:10.1016/S0003-3472(85)80068-3.

Bertrand, V., Bisso, P., Poole, R. J., and Hobert, O. (2011). Notch-dependent induction of left/right asymmetry in C. elegans interneurons and motoneurons. Curr Biol 21, 1225–1231. doi:10.1016/j.cub.2011.06.016.

Bierbach, D., Laskowski, K. L., and Wolf, M. (2017). Behavioural individuality in clonal fish arises despite near-identical rearing conditions. Nat Commun 8, 15361. doi:10.1038/ncomms15361.

Bisazza, A., Pignatti, R., and Vallortigara, G. (1997). Detour tests reveal task-and stimulus-specific behavioural lateralization in mosquitofish (Gambusia holbrooki). Behavioural Brain Research 89, 237–242. doi:10.1016/S0166-4328(97)00061-2.

Blackwood, C. A., Bailetti, A., Nandi, S., Gridley, T., and Hébert, J. M. (2020). Notch Dosage: Jagged1 Haploinsufficiency Is Associated With Reduced Neuronal Division and Disruption of Periglomerular Interneurons in Mice. Front Cell Dev Biol 8, 113. doi:10.3389/fcell.2020.00113.

Brown, C., and Magat, M. (2011). The evolution of lateralized foot use in parrots: a phylogenetic approach. Behavioral Ecology 22, 1201–1208. doi:10.1093/beheco/arr114.

Bruzzone, M., Gatto, E., Lucon Xiccato, T., Dalla Valle, L., Fontana, C. M., Meneghetti, G., et al. (2020). Measuring recognition memory in zebrafish larvae: issues and limitations. PeerJ 8, e8890. doi:10.7717/peerj.8890.

Buchanan, S. M., Kain, J. S., and de Bivort, B. L. (2015). Neuronal control of locomotor handedness in Drosophila. Proc Natl Acad Sci U S A 112, 6700–6705. doi:10.1073/pnas.1500804112.

Bui Quoc, E., Ribot, J., Quenech’Du, N., Doutremer, S., Lebas, N., Grantyn, A., et al. (2012). Asymmetrical Interhemispheric Connections Develop in Cat Visual Cortex after Early Unilateral Convergent Strabismus: Anatomy, Physiology, and Mechanisms. Front. Neuroanat. 0. doi:10.3389/fnana.2011.00068.

Bulman-Fleming, M. B., Bryden, M. P., and Rogers, T. T. (1997). Mouse paw preference: effects of variations in testing protocol. Behav Brain Res 86, 79–87. doi:10.1016/s0166-4328(96)02249-8.

Burgess, H. A., and Granato, M. (2007). Sensorimotor gating in larval zebrafish. J Neurosci 27, 4984–4994. doi:10.1523/JNEUROSCI.0615-07.2007.

Butler, M. G., Iben, J. R., Marsden, K. C., Epstein, J. A., Granato, M., and Weinstein, B. M. (2015). SNPfisher: tools for probing genetic variation in laboratory-reared zebrafish. Development 142, 1542–1552. doi:10.1242/dev.118786.

Byrne, R. A., Kuba, M. J., and Meisel, D. V. (2004). Lateralized eye use in Octopus vulgaris shows antisymmetrical distribution. Animal Behaviour 68, 1107–1114. doi:10.1016/j.anbehav.2003.11.027.

Casey, M. B., and Karpinski, S. (1999). The Development of Postnatal Turning Bias is Influenced by Prenatal Visual Experience in Domestic Chicks (Gallus gallus). Psychol Rec 49, 67–74. doi:10.1007/BF03395307.

Castillo-Ramírez, L. A., Ryu, S., and De Marco, R. J. (2019). Active behaviour during early development shapes glucocorticoid reactivity. Sci Rep 9, 12796. doi:10.1038/s41598-019-49388-3.

Catania, K. C. (2009). Tentacled snakes turn C-starts to their advantage and predict future prey behavior. Proc Natl Acad Sci U S A 106, 11183–11187. doi:10.1073/pnas.0905183106.

Catania, K. C. (2010). Born knowing: tentacled snakes innately predict future prey behavior. PLoS One 5, e10953. doi:10.1371/journal.pone.0010953.

Cauchard, L., Boogert, N. J., Lefebvre, L., Dubois, F., and Doligez, B. (2013). Problem-solving performance is correlated with reproductive success in a wild bird population. Animal Behaviour 85, 19–26. doi:10.1016/j.anbehav.2012.10.005.

Chaumillon, R., Blouin, J., and Guillaume, A. (2018). Interhemispheric Transfer Time Asymmetry of Visual Information Depends on Eye Dominance: An Electrophysiological Study. Front. Neurosci. 0. doi:10.3389/fnins.2018.00072.

Chen, X., and Engert, F. (2014). Navigational strategies underlying phototaxis in larval zebrafish. Front Syst Neurosci 8, 39. doi:10.3389/fnsys.2014.00039.

Chen-Bee, C. H., and Frostig, R. D. (1996). Variability and interhemispheric asymmetry of single-whisker functional representations in rat barrel cortex. Journal of Neurophysiology 76, 884–894. doi:10.1152/jn.1996.76.2.884.

Cheng, Y.-C., Amoyel, M., Qiu, X., Jiang, Y.-J., Xu, Q., and Wilkinson, D. G. (2004). Notch activation regulates the segregation and differentiation of rhombomere boundary cells in the zebrafish hindbrain. Dev Cell 6, 539–550. doi:10.1016/s1534-5807(04)00097-8.

Chu, O., Abeare, C. A., and Bondy, M. A. (2012). Inconsistent vs consistent right-handers’ performance on an episodic memory task: evidence from the California Verbal Learning Test. Laterality 17, 306–317. doi:10.1080/1357650X.2011.568490.

Collins, R. D., Gargesh, R. N., Maltby, A. D., Roggero, R. J., Tourtellot, M. K., and Bell, W. J. (1994). Innate control of local search behaviour in the house fly, Musca domestica. Physiological Entomology 19, 165–172. doi:10.1111/j.1365-3032.1994.tb01039.x.

Collins, R. L. (1969). On the Inheritance of Handedness: II. Selection for sinistrality in mice. Journal of Heredity 60, 117–119. doi:10.1093/oxfordjournals.jhered.a107951.

Coltogirone, R. A., Sherfinski, E. I., Dobler, Z. A., Peterson, S. N., Andlinger, A. R., Fadel, L. C., et al. (2021). Gsx2 but not Gsx1 is necessary for early forebrain patterning and long-term survival in zebrafish. doi:10.1101/2021.09.13.460150.

Colwill, R. M., and Creton, R. (2011). Locomotor behaviors in zebrafish (Danio rerio) larvae. Behav Processes 86, 222–229. doi:10.1016/j.beproc.2010.12.003.

Connor, S. A., Ammendrup-Johnsen, I., Chan, A. W., Kishimoto, Y., Murayama, C., Kurihara, N., et al. (2016). Altered Cortical Dynamics and Cognitive Function upon Haploinsufficiency of the Autism-Linked Excitatory Synaptic Suppressor MDGA2. Neuron 91, 1052–1068. doi:10.1016/j.neuron.2016.08.016.

Conradt, L., Bodsworth, E. J., Roper, T. J., and Thomas, C. D. (2000). Non-random dispersal in the butterfly Maniola jurtina: implications for metapopulation models. Proceedings of the Royal Society of London. Series B: Biological Sciences 267, 1505–1510. doi:10.1098/rspb.2000.1171.

Cuellar-Partida, G., Tung, J. Y., Eriksson, N., Albrecht, E., Aliev, F., Andreassen, O. A., et al. (2020). Genome-wide association study identifies 48 common genetic variants associated with handedness. Nature Human Behaviour, 1–10. doi:10.1038/s41562-020-00956-y.

Davison, J. M., Akitake, C. M., Goll, M. G., Rhee, J. M., Gosse, N., Baier, H., et al. (2007). Transactivation from Gal4-VP16 transgenic insertions for tissue-specific cell labeling and ablation in zebrafish. Dev Biol 304, 811–824. doi:10.1016/j.ydbio.2007.01.033.

De Santi, A., Sovrano, V. A., Bisazza, A., and Vallortigara, G. (2001). Mosquitofish display differential left-and right-eye use during mirror image scrutiny and predator inspection responses. Animal Behaviour 61, 305–310. doi:10.1006/anbe.2000.1566.

De Smedt, M., Hoebeke, I., Reynvoet, K., Leclercq, G., and Plum, J. (2005). Different thresholds of Notch signaling bias human precursor cells toward B-, NK-, monocytic/dendritic-, or T-cell lineage in thymus microenvironment. Blood 106, 3498–3506. doi:10.1182/blood-2005-02-0496.

Demehri, S., Turkoz, A., and Kopan, R. (2009). Epidermal Notch1 loss promotes skin tumorigenesis by impacting the stromal microenvironment. Cancer Cell 16, 55–66. doi:10.1016/j.ccr.2009.05.016.

Derjean, D., Moussaddy, A., Atallah, E., St-Pierre, M., Auclair, F., Chang, S., et al. (2010). A Novel Neural Substrate for the Transformation of Olfactory Inputs into Motor Output. PLOS Biology 8, e1000567. doi:10.1371/journal.pbio.1000567.

Dingemanse, N. J., Both, C., Drent, P. J., and Tinbergen, J. M. (2004). Fitness consequences of avian personalities in a fluctuating environment. Proc Biol Sci 271, 847–852. doi:10.1098/rspb.2004.2680.

Domenici, P., Allan, B. J. M., Watson, S.-A., McCormick, M. I., and Munday, P. L. (2014). Shifting from Right to Left: The Combined Effect of Elevated CO2 and Temperature on Behavioural Lateralization in a Coral Reef Fish. PLOS ONE 9, e87969. doi:10.1371/journal.pone.0087969.

Dreosti, E., Lopes, G., Kampff, A. R., and Wilson, S. W. (2015). Development of social behavior in young zebrafish. Front Neural Circuits 9, 39. doi:10.3389/fncir.2015.00039.

Dreosti, E., Vendrell Llopis, N., Carl, M., Yaksi, E., and Wilson, S. W. (2014). Left-right asymmetry is required for the habenulae to respond to both visual and olfactory stimuli. Curr. Biol. 24, 440–445. doi:10.1016/j.cub.2014.01.016.

Dunn, T. W., Mu, Y., Narayan, S., Randlett, O., Naumann, E. A., Yang, C.-T., et al. (2016). Brain-wide mapping of neural activity controlling zebrafish exploratory locomotion. Elife 5, e12741. doi:10.7554/eLife.12741.

Elnitsky, M. A., and Claussen, D. L. (2006). The effects of temperature and inter-individual variation on the locomotor performance of juvenile turtles. J Comp Physiol B 176, 497–504. doi:10.1007/s00360-006-0071-1.

Engeszer, R. E., Patterson, L. B., Rao, A. A., and Parichy, D. M. (2007). Zebrafish in The Wild: A Review of Natural History And New Notes from The Field. Zebrafish 4, 21–40. doi:10.1089/zeb.2006.9997.

Eto, K., Mazilu-Brown, J. K., Henderson-MacLennan, N., Dipple, K. M., and McCabe, E. R. B. (2014). Development of catecholamine and cortisol stress responses in zebrafish. Molecular Genetics and Metabolism Reports 1, 373–377. doi:10.1016/j.ymgmr.2014.08.003.

Fauq, A. H., Simpson, K., Maharvi, G. M., Golde, T., and Das, P. (2007). A multigram chemical synthesis of the gamma-secretase inhibitor LY411575 and its diastereoisomers. Bioorg Med Chem Lett 17, 6392–6395. doi:10.1016/j.bmcl.2007.07.062.

Fernandes, A. M., Mearns, D. S., Donovan, J. C., Larsch, J., Helmbrecht, T. O., Kölsch, Y., et al. (2021). Neural circuitry for stimulus selection in the zebrafish visual system. Neuron 109, 805–822.e6. doi:10.1016/j.neuron.2020.12.002.

Filosa, A., Barker, A. J., Dal Maschio, M., and Baier, H. (2016). Feeding State Modulates Behavioral Choice and Processing of Prey Stimuli in the Zebrafish Tectum. Neuron 90, 596–608. doi:10.1016/j.neuron.2016.03.014.

Fischer-Zirnsak, B., Segebrecht, L., Schubach, M., Charles, P., Alderman, E., Brown, K., et al. (2019). Haploinsufficiency of the Notch Ligand DLL1 Causes Variable Neurodevelopmental Disorders. The American Journal of Human Genetics 105, 631–639. doi:10.1016/j.ajhg.2019.07.002.

Fontana, B. D., Cleal, M., Clay, J. M., and Parker, M. O. (2019). Zebrafish (Danio rerio) behavioral laterality predicts increased short-term avoidance memory but not stress-reactivity responses. Anim Cogn 22, 1051–1061. doi:10.1007/s10071-019-01296-9.

Freund, J., Brandmaier, A. M., Lewejohann, L., Kirste, I., Kritzler, M., Krüger, A., et al. (2013). Emergence of individuality in genetically identical mice. Science 340, 756–759. doi:10.1126/science.1235294.

Freund, J., Brandmaier, A. M., Lewejohann, L., Kirste, I., Kritzler, M., Krüger, A., et al. (2015). Association between exploratory activity and social individuality in genetically identical mice living in the same enriched environment. Neuroscience 309, 140–152. doi:10.1016/j.neuroscience.2015.05.027.

Freund, N., Valencia-Alfonso, C. E., Kirsch, J., Brodmann, K., Manns, M., and Güntürkün, O. (2016). Asymmetric top-down modulation of ascending visual pathways in pigeons. Neuropsychologia 83, 37–47. doi:10.1016/j.neuropsychologia.2015.08.014.

Gahtan, E., Tanger, P., and Baier, H. (2005). Visual Prey Capture in Larval Zebrafish Is Controlled by Identified Reticulospinal Neurons Downstream of the Tectum. J. Neurosci. 25, 9294–9303. doi:10.1523/JNEUROSCI.2678-05.2005.

Gamse, J. T., Kuan, Y.-S., Macurak, M., Brösamle, C., Thisse, B., Thisse, C., et al. (2005). Directional asymmetry of the zebrafish epithalamus guides dorsoventral innervation of the midbrain target. Development 132, 4869–4881. doi:10.1242/dev.02046.

Gamse, J. T., Thisse, C., Thisse, B., and Halpern, M. E. (2003). The parapineal mediates left-right asymmetry in the zebrafish diencephalon. Development 130, 1059–1068. doi:10.1242/dev.00270.

Gebhardt, C., Auer, T. O., Henriques, P. M., Rajan, G., Duroure, K., Bianco, I. H., et al. (2019). An interhemispheric neural circuit allowing binocular integration in the optic tectum. Nat Commun 10, 5471. doi:10.1038/s41467-019-13484-9.

Geling, A., Steiner, H., Willem, M., Bally-Cuif, L., and Haass, C. (2002). A γ-secretase inhibitor blocks Notch signaling in vivo and causes a severe neurogenic phenotype in zebrafish. EMBO Rep 3, 688–694. doi:10.1093/embo-reports/kvf124.

Gibbons, T. C., Rudman, S. M., and Schulte, P. M. (2017). Low temperature and low salinity drive putatively adaptive growth differences in populations of threespine stickleback. Sci Rep 7, 16766. doi:10.1038/s41598-017-16919-9.

Giljov, A., Karenina, K., and Malashichev, Y. (2013). Forelimb preferences in quadrupedal marsupials and their implications for laterality evolution in mammals. BMC Evol Biol 13, 61. doi:10.1186/1471-2148-13-61.

Gray, J. M., Hill, J. J., and Bargmann, C. I. (2005). A circuit for navigation in Caenorhabditis elegans. Proc Natl Acad Sci U S A 102, 3184–3191. doi:10.1073/pnas.0409009101.

Güntürkün, O., Hellmann, B., Melsbach, G., and Prior, H. (1998). Asymmetries of representation in the visual system of pigeons. Neuroreport 9, 4127–4130. doi:10.1097/00001756-199812210-00023.

Haesemeyer, M., Robson, D. N., Li, J. M., Schier, A. F., and Engert, F. (2018). A Brain-wide Circuit Model of Heat-Evoked Swimming Behavior in Larval Zebrafish. Neuron 98, 817–831.e6. doi:10.1016/j.neuron.2018.04.013.

Hager, T., Jansen, R. F., Pieneman, A. W., Manivannan, S. N., Golani, I., van der Sluis, S., et al. (2014). Display of individuality in avoidance behavior and risk assessment of inbred mice. Front. Behav. Neurosci. 0. doi:10.3389/fnbeh.2014.00314.

Hills, T., Brockie, P. J., and Maricq, A. V. (2004). Dopamine and Glutamate Control Area-Restricted Search Behavior in Caenorhabditis elegans. J Neurosci 24, 1217–1225. doi:10.1523/JNEUROSCI.1569-03.2004.

Hills, T. T., Kalff, C., and Wiener, J. M. (2013). Adaptive Lévy Processes and Area-Restricted Search in Human Foraging. PLOS ONE 8, e60488. doi:10.1371/journal.pone.0060488.

Horstick, E. J., Bayleyen, Y., and Burgess, H. A. (2020). Molecular and cellular determinants of motor asymmetry in zebrafish. Nat Commun 11, 1170. doi:10.1038/s41467-020-14965-y.

Horstick, E. J., Bayleyen, Y., Sinclair, J. L., and Burgess, H. A. (2017). Search strategy is regulated by somatostatin signaling and deep brain photoreceptors in zebrafish. BMC Biol. 15, 4. doi:10.1186/s12915-016-0346-2.

Horstick, E. J., Linsley, J. W., Dowling, J. J., Hauser, M. A., McDonald, K. K., Ashley-Koch, A., et al. (2013). Stac3 is a component of the excitation-contraction coupling machinery and mutated in Native American myopathy. Nat Commun 4, 1952. doi:10.1038/ncomms2952.

Horváth, G., Jiménez-Robles, O., Martín, J., López, P., De la Riva, I., and Herczeg, G. (2020). Linking behavioral thermoregulation, boldness, and individual state in male Carpetan rock lizards. Ecol Evol 10, 10230–10241. doi:10.1002/ece3.6685.

Humphries, D. A., and Driver, P. M. (1970). Protean defence by prey animals. Oecologia 5, 285–302. doi:10.1007/BF00815496.

Ishii, K., Wohl, M., DeSouza, A., and Asahina, K. (2020). Sex-determining genes distinctly regulate courtship capability and target preference via sexually dimorphic neurons. Elife 9, e52701. doi:10.7554/eLife.52701.

Itoh, M., Kim, C.-H., Palardy, G., Oda, T., Jiang, Y.-J., Maust, D., et al. (2003). Mind bomb is a ubiquitin ligase that is essential for efficient activation of Notch signaling by Delta. Dev Cell 4, 67–82. doi:10.1016/s1534-5807(02)00409-4.

Izvekov, E. I., Nepomnyashchikh, V. A., Medyantseva, E. N., Chebotareva, Y. V., and Izyumov, Y. G. (2012). Selection of direction of movement and bilateral morphological asymmetry in young roach (Rutilus rutilus). Biol Bull Rev 2, 364–370. doi:10.1134/S2079086412040044.

Jacobs, C. T., and Huang, P. (2019). Notch signalling maintains Hedgehog responsiveness via a Gli-dependent mechanism during spinal cord patterning in zebrafish. eLife 8. doi:10.7554/eLife.49252.

Jäncke, L., and Steinmetz, H. (1995). Hand motor performance and degree of asymmetry in monozygotic twins. Cortex 31, 779–785. doi:10.1016/s0010-9452(13)80028-7.

Kain, J. S., Stokes, C., and de Bivort, B. L. (2012). Phototactic personality in fruit flies and its suppression by serotonin and white. Proc Natl Acad Sci U S A 109, 19834–19839. doi:10.1073/pnas.1211988109.

Kain, J. S., Zhang, S., Akhund-Zade, J., Samuel, A. D. T., Klein, M., and de Bivort, B. L. (2015). Variability in thermal and phototactic preferences in Drosophila may reflect an adaptive bet-hedging strategy. Evolution 69, 3171–3185. doi:10.1111/evo.12813.

Kim, C. H., Ueshima, E., Muraoka, O., Tanaka, H., Yeo, S. Y., Huh, T. L., et al. (1996). Zebrafish elav/HuC homologue as a very early neuronal marker. Neurosci Lett 216, 109–112. doi:10.1016/0304-3940(96)13021-4.

Kimmel, C. B., Ballard, W. W., Kimmel, S. R., Ullmann, B., and Schilling, T. F. (1995). Stages of embryonic development of the zebrafish. Dev Dyn 203, 253–310. doi:10.1002/aja.1002030302.

Klein, S., Pasquaretta, C., Barron, A. B., Devaud, J.-M., and Lihoreau, M. (2017). Inter-individual variability in the foraging behaviour of traplining bumblebees. Sci Rep 7, 4561. doi:10.1038/s41598-017-04919-8.

Körholz, J. C., Zocher, S., Grzyb, A. N., Morisse, B., Poetzsch, A., Ehret, F., et al. (2018). Selective increases in inter-individual variability in response to environmental enrichment in female mice. eLife 7, e35690. doi:10.7554/eLife.35690.

Krebs, L. T., Shutter, J. R., Tanigaki, K., Honjo, T., Stark, K. L., and Gridley, T. (2004). Haploinsufficient lethality and formation of arteriovenous malformations in Notch pathway mutants. Genes Dev 18, 2469–2473. doi:10.1101/gad.1239204.

Kumar, A., Huh, T.-L., Choe, J., and Rhee, M. (2017). Rnf152 Is Essential for NeuroD Expression and Delta-Notch Signaling in the Zebrafish Embryos. Mol Cells 40, 945–953. doi:10.14348/molcells.2017.0216.

Kushlan, J. A. (1976). Feeding behavior of North Americans herons. The Auk 93, 86–94.

LaFoya, B., Munroe, J. A., Mia, M. M., Detweiler, M. A., Crow, J. J., Wood, T., et al. (2016). Notch: A multi-functional integrating system of microenvironmental signals. Dev Biol 418, 227–241. doi:10.1016/j.ydbio.2016.08.023.

Lambert, A. M., Bonkowsky, J. L., and Masino, M. A. (2012). The Conserved Dopaminergic Diencephalospinal Tract Mediates Vertebrate Locomotor Development in Zebrafish Larvae. J. Neurosci. 32, 13488–13500. doi:10.1523/JNEUROSCI.1638-12.2012.

Larsch, J., and Baier, H. (2018). Biological Motion as an Innate Perceptual Mechanism Driving Social Affiliation. Current Biology 28, 3523–3532.e4. doi:10.1016/j.cub.2018.09.014.

Lewis, L. P. C., Siju, K. P., Aso, Y., Friedrich, A. B., Bulteel, A. J. B., Rubin, G. M., et al. (2015). A Higher Brain Circuit for Immediate Integration of Conflicting Sensory Information in Drosophila. Curr Biol 25, 2203–2214. doi:10.1016/j.cub.2015.07.015.

Linneweber, G. A., Andriatsilavo, M., Dutta, S. B., Bengochea, M., Hellbruegge, L., Liu, G., et al. (2020). A neurodevelopmental origin of behavioral individuality in the Drosophila visual system. Science 367, 1112–1119. doi:10.1126/science.aaw7182.

Long, Y., Li, L., Li, Q., He, X., and Cui, Z. (2012). Transcriptomic Characterization of Temperature Stress Responses in Larval Zebrafish. PLOS ONE 7, e37209. doi:10.1371/journal.pone.0037209.

Manns, M., Otto, T., and Salm, L. (2021). Pigeons show how meta-control enables decision-making in an ambiguous world. Sci Rep 11, 3838. doi:10.1038/s41598-021-83406-7.

Manns, M., and Römling, J. (2012). The impact of asymmetrical light input on cerebral hemispheric specialization and interhemispheric cooperation. Nat Commun 3, 696. doi:10.1038/ncomms1699.

Marquart, G. D., Tabor, K. M., Bergeron, S. A., Briggman, K. L., and Burgess, H. A. (2019). Prepontine non-giant neurons drive flexible escape behavior in zebrafish. PLoS Biol. 17, e3000480. doi:10.1371/journal.pbio.3000480.

Marquart, G. D., Tabor, K. M., Brown, M., Strykowski, J. L., Varshney, G. K., LaFave, M. C., et al. (2015). A 3D Searchable Database of Transgenic Zebrafish Gal4 and Cre Lines for Functional Neuroanatomy Studies. Front. Neural Circuits 9. doi:10.3389/fncir.2015.00078.

Matsuda, M., Rand, K., Palardy, G., Shimizu, N., Ikeda, H., Dalle Nogare, D., et al. (2016). Epb41l5 competes with Delta as a substrate for Mib1 to coordinate specification and differentiation of neurons. Development 143, 3085–3096. doi:10.1242/dev.138743.

Miyashita, T., and Palmer, A. R. (2014). Handed behavior in hagfish--an ancient vertebrate lineage--and a survey of lateralized behaviors in other invertebrate chordates and elongate vertebrates. Biol Bull 226, 111–120. doi:10.1086/BBLv226n2p111.

Mizutani, K., Yoon, K., Dang, L., Tokunaga, A., and Gaiano, N. (2007). Differential Notch signalling distinguishes neural stem cells from intermediate progenitors. Nature 449, 351–355. doi:10.1038/nature06090.

Muto, A., Lal, P., Ailani, D., Abe, G., Itoh, M., and Kawakami, K. (2017). Activation of the hypothalamic feeding centre upon visual prey detection. Nat Commun 8, 15029. doi:10.1038/ncomms15029.

Nelson, J. C., Witze, E., Ma, Z., Ciocco, F., Frerotte, A., Randlett, O., et al. (2020). Acute Regulation of Habituation Learning via Posttranslational Palmitoylation. Curr Biol 30, 2729–2738.e4. doi:10.1016/j.cub.2020.05.016.

Ohata, S., Aoki, R., Kinoshita, S., Yamaguchi, M., Tsuruoka-Kinoshita, S., Tanaka, H., et al. (2011). Dual roles of Notch in regulation of apically restricted mitosis and apicobasal polarity of neuroepithelial cells. Neuron 69, 215–230. doi:10.1016/j.neuron.2010.12.026.

Pantoja, C., Hoagland, A., Carroll, E. C., Karalis, V., Conner, A., and Isacoff, E. Y. (2016). Neuromodulatory Regulation of Behavioral Individuality in Zebrafish. Neuron 91, 587–601. doi:10.1016/j.neuron.2016.06.016.

Pantoja, C., Larsch, J., Laurell, E., Marquart, G., Kunst, M., and Baier, H. (2020). Rapid Effects of Selection on Brain-wide Activity and Behavior. Current Biology 30, 3647–3656.e3. doi:10.1016/j.cub.2020.06.086.

Pei, Z., Wang, B., Chen, G., Nagao, M., Nakafuku, M., and Campbell, K. (2011). Homeobox genes Gsx1 and Gsx2 differentially regulate telencephalic progenitor maturation. Proc Natl Acad Sci U S A 108, 1675–1680. doi:10.1073/pnas.1008824108.

R Core Team (2020). R: A language and environment for statistical computing (2020). Vienna, Austria Available at: https://www.R-project.org/.

Rogers, L. J. (1982). Light experience and asymmetry of brain function in chickens. Nature 297, 223–225. doi:10.1038/297223a0.

Rogers, L. J., and Deng, C. (2005). Corticosterone treatment of the chick embryo affects light-stimulated development of the thalamofugal visual pathway. Behav. Brain Res. 159, 63–71. doi:10.1016/j.bbr.2004.10.003.

Roussigné, M., Bianco, I. H., Wilson, S. W., and Blader, P. (2009). Nodal signalling imposes left-right asymmetry upon neurogenesis in the habenular nuclei. Development 136, 1549–1557. doi:10.1242/dev.034793.

Roychoudhury, K., Salomone, J., Qin, S., Cain, B., Adam, M., Potter, S. S., et al. (2020). Physical interactions between Gsx2 and Ascl1 balance progenitor expansion versus neurogenesis in the mouse lateral ganglionic eminence. Development 147, dev185348. doi:10.1242/dev.185348.

Schiffner, I., and Srinivasan, M. V. (2013). Behavioural lateralization in Budgerigars varies with the task and the individual. PLoS One 8, e82670. doi:10.1371/journal.pone.0082670.

Schwarzkopf, M., Liu, M. C., Schulte, S. J., Ives, R., Husain, N., Choi, H. M. T., et al. (2021). Hybridization chain reaction enables a unified approach to multiplexed, quantitative, high-resolution immunohistochemistry and in situ hybridization. bioRxiv, 2021.06.02.446311. doi:10.1101/2021.06.02.446311.

Sharma, P., Saraswathy, V. M., Xiang, L., and Fürthauer, M. (2019). Notch-mediated inhibition of neurogenesis is required for zebrafish spinal cord morphogenesis. Sci Rep 9, 9958. doi:10.1038/s41598-019-46067-1.

Shen, W., Huang, J., and Wang, Y. (2021). Biological Significance of NOTCH Signaling Strength. Frontiers in Cell and Developmental Biology 9, 570. doi:10.3389/fcell.2021.652273.

Simons, A. M. (2011). Modes of response to environmental change and the elusive empirical evidence for bet hedging. Proceedings of the Royal Society B: Biological Sciences 278, 1601–1609. doi:10.1098/rspb.2011.0176.

Souman, J. L., Frissen, I., Sreenivasa, M. N., and Ernst, M. O. (2009). Walking straight into circles. Curr Biol 19, 1538–1542. doi:10.1016/j.cub.2009.07.053.

Sovrano, V. A. (2004). Visual lateralization in response to familiar and unfamiliar stimuli in fish. Behavioural Brain Research 152, 385–391. doi:10.1016/j.bbr.2003.10.022.

Stern, S., Kirst, C., and Bargmann, C. I. (2017). Neuromodulatory Control of Long-Term Behavioral Patterns and Individuality across Development. Cell 171, 1649–1662.e10. doi:10.1016/j.cell.2017.10.041.

Striedter, G. F. (1991). Auditory, electrosensory, and mechanosensory lateral line pathways through the forebrain in channel catfishes. Journal of Comparative Neurology 312, 311–331. doi:10.1002/cne.903120213.

Sundin, J., Morgan, R., Finnøen, M. H., Dey, A., Sarkar, K., and Jutfelt, F. (2019). On the Observation of Wild Zebrafish (Danio rerio) in India. Zebrafish 16, 546–553. doi:10.1089/zeb.2019.1778.

Tallafuss, A., Trepman, A., and Eisen, J. S. (2009). DeltaA mRNA and protein distribution in the zebrafish nervous system. Dev Dyn 238, 3226–3236. doi:10.1002/dvdy.22136.

Tremblay, Y., Roberts, A. J., and Costa, D. P. (2007). Fractal landscape method: an alternative approach to measuring area-restricted searching behavior. J Exp Biol 210, 935–945. doi:10.1242/jeb.02710.

Versace, E., Caffini, M., Werkhoven, Z., and de Bivort, B. L. (2020). Individual, but not population asymmetries, are modulated by social environment and genotype in Drosophila melanogaster. Sci Rep 10, 4480. doi:10.1038/s41598-020-61410-7.

Vogt, G., Huber, M., Thiemann, M., Boogaart, G. van den, Schmitz, O. J., and Schubart, C. D. (2008). Production of different phenotypes from the same genotype in the same environment by developmental variation. Journal of Experimental Biology 211, 510–523. doi:10.1242/jeb.008755.

Wang, B., Waclaw, R. R., Allen, Z. J., Guillemot, F., and Campbell, K. (2009). Ascl1 is a required downstream effector of Gsx gene function in the embryonic mouse telencephalon. Neural Dev 4, 5. doi:10.1186/1749-8104-4-5.

Watson, N. V., and Kimura, D. (1989). Right-hand superiority for throwing but not for intercepting. Neuropsychologia 27, 1399–1414. doi:10.1016/0028-3932(89)90133-4.

Wickham, H. (2016). ggplot2: Elegant Graphics for Data Analysis. Springer-Verlag New York.

Yaeger, C., Ros, A. M., Cross, V., DeAngelis, R. S., Stobaugh, D. J., and Rhodes, J. S. (2014). Blockade of arginine vasotocin signaling reduces aggressive behavior and c-Fos expression in the preoptic area and periventricular nucleus of the posterior tuberculum in male Amphiprion ocellaris. Neuroscience 267, 205–218. doi:10.1016/j.neuroscience.2014.02.045.

Yapici, N., Cohn, R., Schusterreiter, C., Ruta, V., and Vosshall, L. B. (2016). A Taste Circuit that Regulates Ingestion by Integrating Food and Hunger Signals. Cell 165, 715–729. doi:10.1016/j.cell.2016.02.061.

Yokogawa, T., Hannan, M. C., and Burgess, H. A. (2012). The Dorsal Raphe Modulates Sensory Responsiveness during Arousal in Zebrafish. J. Neurosci. 32, 15205–15215. doi:10.1523/JNEUROSCI.1019-12.2012.

Yoon, K.-J., Koo, B.-K., Im, S.-K., Jeong, H.-W., Ghim, J., Kwon, M.-C., et al. (2008). Mind bomb 1-expressing intermediate progenitors generate notch signaling to maintain radial glial cells. Neuron 58, 519–531. doi:10.1016/j.neuron.2008.03.018.

Zadicario, P., Avni, R., Zadicario, E., and Eilam, D. (2005). ‘Looping’-an exploration mechanism in a dark open field. Behav Brain Res 159, 27–36. doi:10.1016/j.bbr.2004.09.022.

Zocher, S., Schilling, S., Grzyb, A. N., Adusumilli, V. S., Bogado Lopes, J., Günther, S., et al. (2020). Early-life environmental enrichment generates persistent individualized behavior in mice. Sci Adv 6, eabb1478. doi:10.1126/sciadv.abb1478.

