## Supplementary Figures for "Environmental and molecular modulation of motor individuality in larval zebrafish"

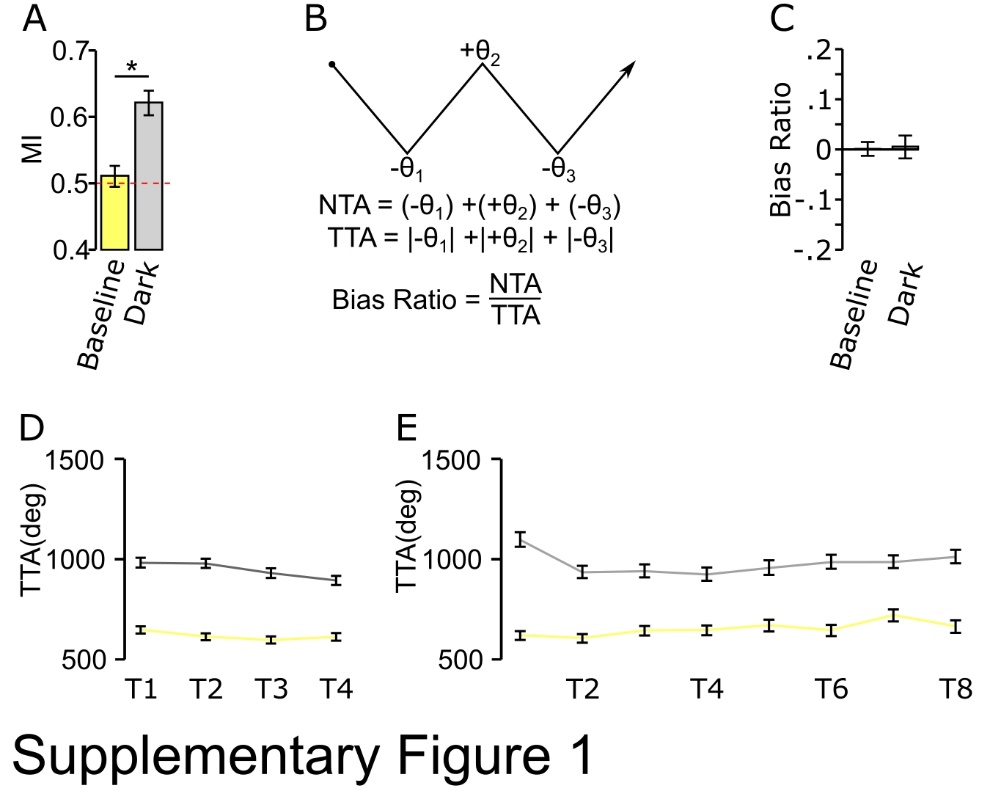

**Supplemental Figure 1. (A)** 4X match index for paired baseline (yellow) and dark response (grey). Red dotted line indicates random choice. Asterisk, p <0.05 Wilcoxon signed-rank test (N = 374). **(B)** Representation and calculation used for generating bias ratio. NTA is the sum of left (-) and right (+) angular displacement, while TTA is the absolute sum of all angular displacement. Bias ratio is the dividend of NTA to TTA. **(C)** Population average of 4X recordings (n = 374) **(D-E)** 4x (N=374) and 8X (N=189) TTA for baseline (yellow) and dark response (grey).

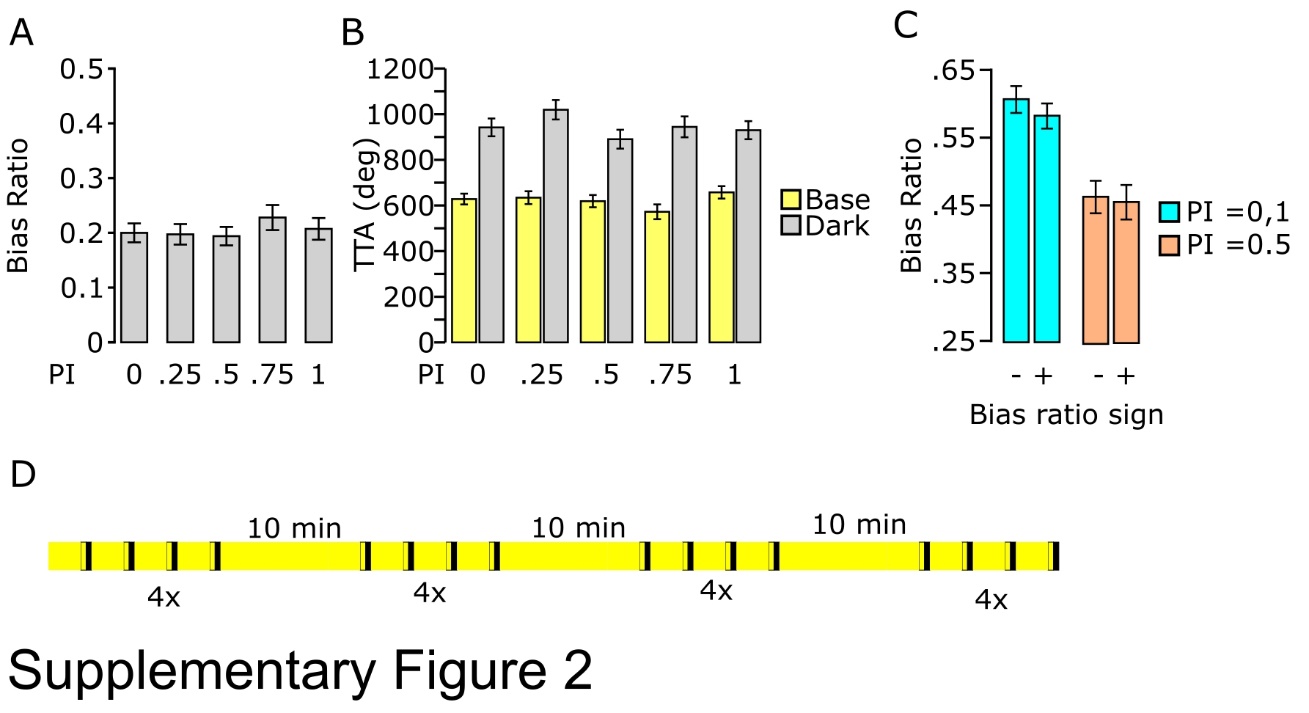

**Supplemental Figure 2. (A)** Absolute bias ratio for 4x baseline illumination responses per PI (PI 0, N=66; PI 0.25, N =74; PI 0.5, N=75; PI 0.75, N=75; PI 1, N= 67). **(B)** TTA for data in A, showing baseline (yellow) and dark (grey) responses. **(C)** Single event absolute bias ratio for matched (PI 0,1 cyan) and unbiased (PI 0.5, orange) individuals sorted by direction (+, rightward; -, leftward). **(D)** Diagram of q4x recording. Yellow and black indicating lights on and off, respectively. Black outline areas indicate 30 second recording windows.

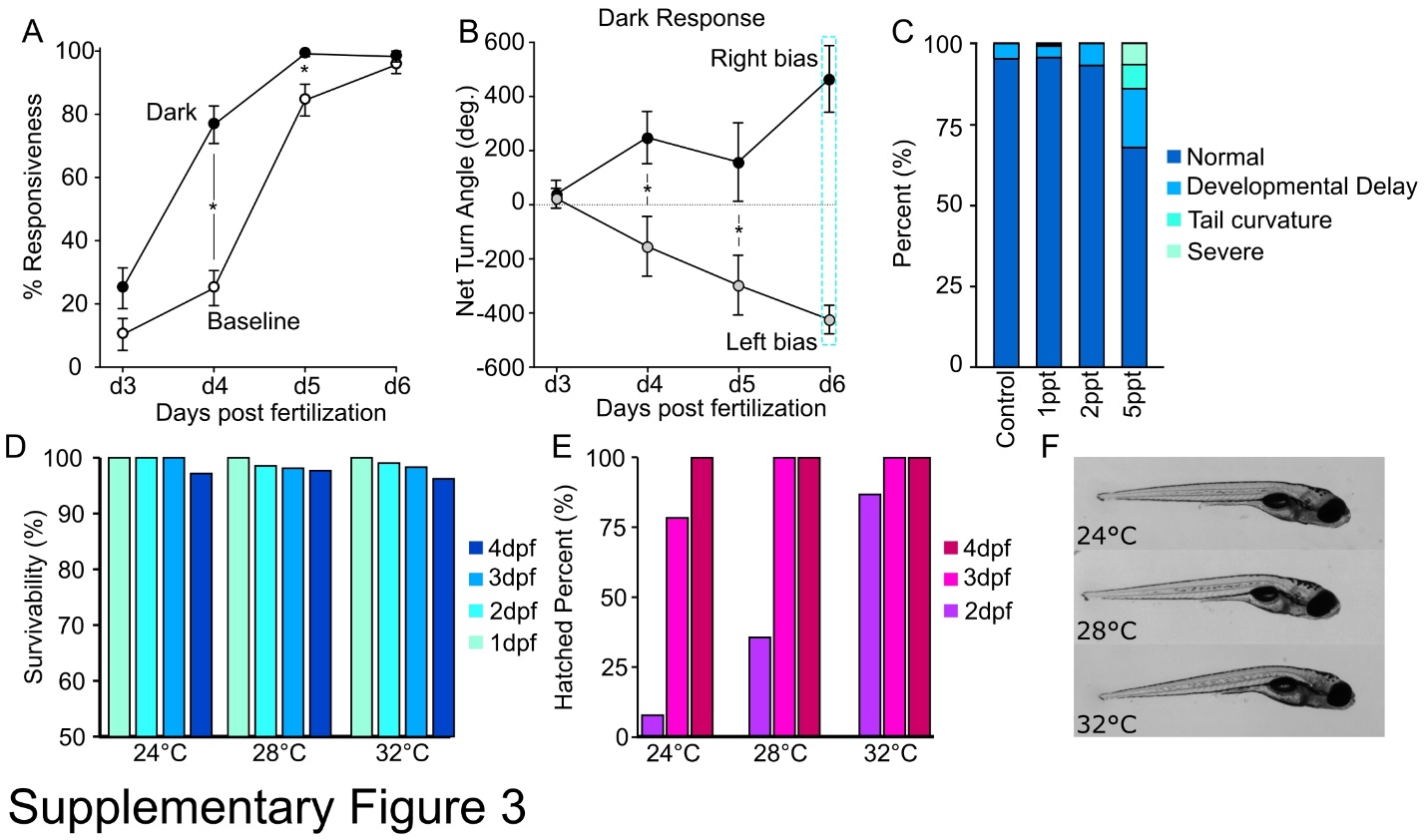

**Supplemental Figure 3. (A-B)** Developmental onset of turn bias. Individual larvae were tested daily in a 4X assay from 3 to 6 dpf. **(A)** Proportion of individuals that show motor responsiveness (N=29) **(B)** Presence of turn bias during development. At 6dpf (dotted blue box), larvae were categorized as right (N=11) or left (N=18) bias based on average NTA (+, right bias; -, left bias) to group responses over the testing interval. Asterisk p <0.05, *t*-test between points at same developmental stage in A and B. No significance shown for 6dpf in B as this timepoint was used to group larvae by performance. **(C)** Phenotypic scoring at 4dpf following variable salinity exposures during early development. As 5ppt generated some developmental abnormalities this treatment was excluded from behavior testing. **(D)** Survival of larvae raised from 1-4dpf at different temperatures (24°C, N=103; 28°C, N=180; 32°C, N=209). **(E)** Proportion of hatched (dechorinated) embryos due to different temperatures (24°C, N=103; 28°C, N=180; 32°C, N=209). **(F)** Representative images of 6dpf larvae after varying temperature exposure from 1-4 dpf.

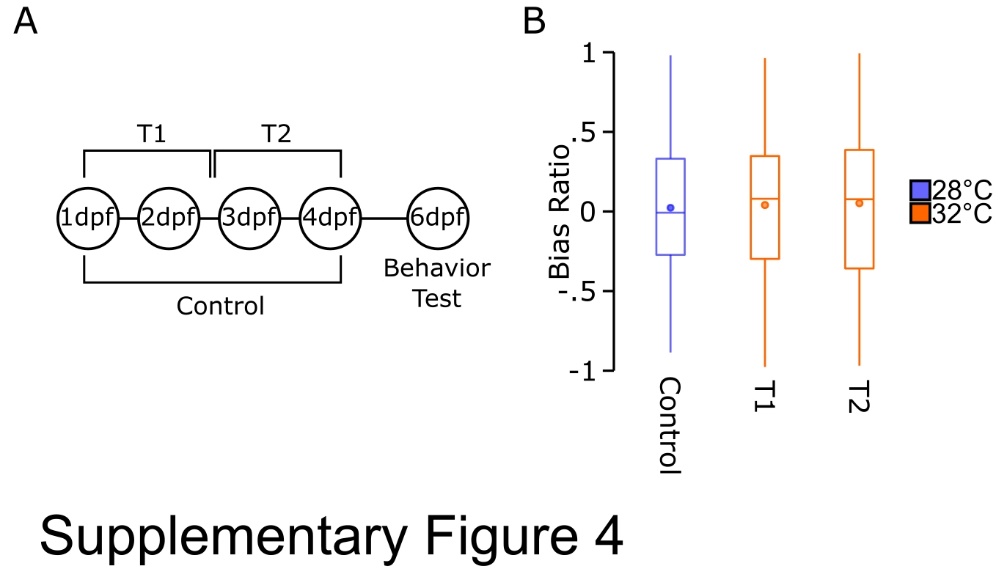

**Supplemental Figure 4. (A)** Timeline of elevated temperature testing showing control (28°C) and T1 (31-55 hpf, 32°C) and T2 (55-79 hpf, 32°C) testing conditions which are not statistically different from random (1 sample t-test against 0: control- 0.023 ± 0.043 *t(100)=0.53*, p=0.60; T1- 0.040 ± 0.042 *t(100)=0.97*, p=0.33; T2- 0.052 ± 0.045 *t(106)=1.151*, p=0.25). **(B)** Average population bias ratio for 28°C control (purple, N=101) and T1 (orange, N=111) and T2 (orange, N=107) elevated temperature groups. Circles show means.

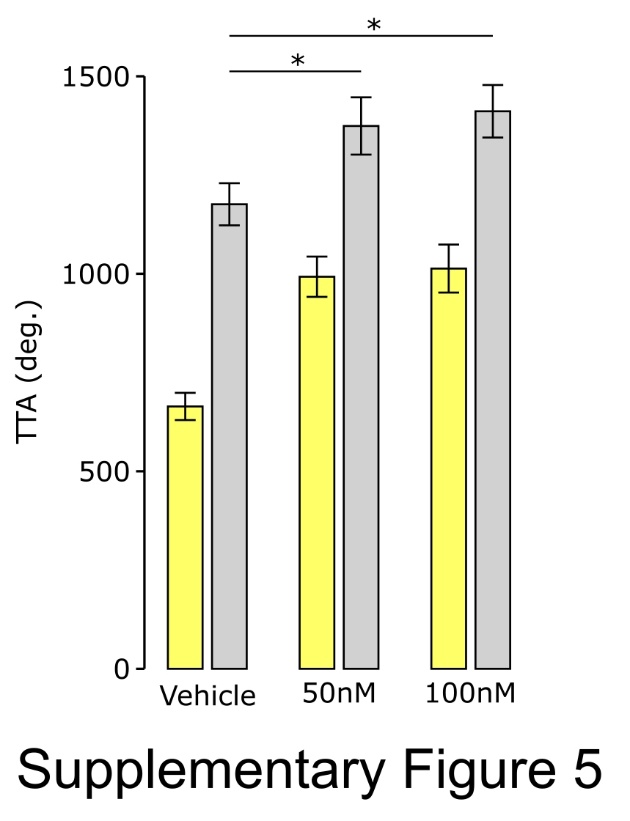

**Supplemental Figure 5**. TTA following Notch inhibitor exposure during early development (vehicle, N=69; 50µM, N=69; 100µM, N=48).
